# Shivering thermogenesis driven by hypothermia-sensitive neurons in the dorsomedial hypothalamus

**DOI:** 10.1101/2025.11.27.690920

**Authors:** Shuai Zhang, Han Zhang, Zitian Huang, Shiyu Wu, Xinyi Yang, Wei He, Donghai Li, Zhenyu Zhang, Yunxuan Wang, Lixin Huang, Runbo Liu, Shuifa Chen, Sheng-Tao Hou

## Abstract

Hypothermia, a decline in core body temperature below 35°C in homeotherms, elicits rapid, asynchronous involuntary shivering thermogenesis; however, the hypothalamic neurons responsible for initiating this response remain uncharacterized. Here, retrograde transsynaptic tracing combined with RNA-seq identified a distinct population of VGLUT2-positive neurons enriched with Calbindin D28k (CALB1) within the dorsomedial hypothalamic nucleus (DMH^Calb1/Vglut2^). Convergent evidence from chemogenetic, optogenetic, fiber photometry calcium imaging, and electromyography (EMG) experiments demonstrated that these neurons selectively mediate hypothermia-triggered shivering thermogenesis. Bidirectional perturbation of DMH^Calb1/Vglut2^ activity modulated shivering intensity, while CALB1-negative VGLUT2-expressing DMH neurons were responsible for non-shivering thermogenesis (NST). Genetic ablation of afferent inputs from LSI/POA/VLPAG projecting to DMH^Calb1/Vglut2^, as well as *ex vivo* pharmacological synaptic blockade, failed to eliminate shivering responses during hypothermia, indicating that their activation is independent of canonical cold-responsive circuits. Hypothermia significantly suppressed transient A-type K⁺ (*I*_A_) currents, without affecting delayed rectifier K⁺ (*I*_K_) currents, accompanied by a pronounced reduction in the *I*_A_ inactivation time constant relative to normothermic 36 °C. Collectively, hypothermia elevates DMH^Calb1/Vglut2^ excitability through *I*_A_-dependent reshaping of action potential dynamics. Shortened action potential repolarization and afterhyperpolarization duration cooperatively boost neuronal firing, constituting an intrinsic mechanism for DMH^Calb1/Vglut2^ neuron-driven shivering initiation.

## Introduction

Homeothermic mammals maintain core body temperature (T_core_) within a narrow homeostatic range of 36-37°C to preserve metabolic and physiological stability ^1, 2^. This process is governed by the hypothalamus, a central thermostat that integrates afferent inputs from peripheral thermoreceptors to coordinate downstream autonomic thermoeffector pathways ^1, 3–5^, balancing heat dissipation and conservation across fluctuating ambient temperatures.

Cold exposure activates two primary thermogenic pathways: non-shivering thermogenesis (NST) and shivering thermogenesis ^6–10^. While NST coordinates metabolic and cardiovascular adaptations, such as tachycardia, vasodilation, elevated brown adipose tissue thermogenesis (T_BAT_), and increased locomotion ^5, 11–13^, shivering serves as the primary defense against cold-mediated heat loss, generating up to 60% of required thermogenesis through skeletal muscle contractions ^14^.

Thermoregulatory muscle responses change dynamically based on cold severity. Initial drops in skin temperature elevate muscle tone, a subtle, isometric “pre-shivering” contraction of the muscles ^15–17^. This sustained tension recruits fatigue-resistant, slow-twitch (Type I) muscle fibers to produce heat without causing overt tremor or shivering ^15, 17^. Additionally, muscle tone also plays a major role in maintaining posture, stabilizing joints, and acting during pain ^18^. Conversely, severe or sustained declines in ambient or T_core_, such as during hypothermia when T_core_ falls below 35°C in humans, trigger visible shivering, characterized by rapid, asynchronous, and involuntary bursts of muscle contractions. This high-potency mechanism recruits powerful, fatigue-prone, fast-twitch (Type II) fibers, increasing metabolic heat production up to 6-fold above basal levels ^15, 17, 19^. It is now recognized that muscle tone and overt shivering are markedly distinct phenomena underlying muscle-driven thermogenesis ^17^, the former responds to ambient temperature fluctuations above 7°C to maintain normothermia, while the latter is triggered during hypothermia when T_core_ drops below 35°C.

However, the exact neural control mechanisms of hypothermia-evoked shivering thermogenesis remain poorly understood. Early investigations established the necessity of the anterior and posterior hypothalamus in regulating shivering behavior ^13, 20, 21^. More recent mapping has delineated a core ascending and descending neurocircuitry encompassing the preoptic area (POA), median preoptic nucleus (MPN), dorsomedial hypothalamus (DMH), and rostral raphe pallidus (rRPa) ^2, 3, 9, 11, 21–23^. Within this circuit, cutaneous cold signals are relayed via the POA to the DMH, which ultimately drives rRPa neurons to activate downstream motor units and skeletal muscle thermogenic responses ^1, 6, 9, 24, 25^. However, this cold-sensing circuit was validated under conditions in which the ambient temperature decreased slowly, without ever falling below 7°C, while the T_core_ remained within the normothermic range. No visible phasic burst shivering occurs under this condition. We have shown that lowering T_core_ to 35-33°C in freely behaving mice triggers strong overt burst shivering ^26^. It is still unclear whether the canonical cold-responsive signaling pathway also drives hypothermia-evoked shivering thermogenesis.

Nevertheless, specific hypothalamic neurons have been identified as key regulators of NST and associated autonomic outputs, including T_BAT_, white adipose browning, heart rate, and blood pressure. These neurons include dmVMH^Pdyn^ excitatory neurons ^6^, the POA^BRS3^ neurons ^27^, and DMH^GABA^ neurons ^28^. Contrary to the common belief that cold-sensitive hypothalamic neurons are rare, Kobayashi (1986) showed that these neurons actually remain electrically silent at a murine normothermic T_core_ of approximately 38 °C ^29^. These neurons fire only at hypothermic temperatures of 36-32LJ°C and display threshold temperature responses, and transient temperature-dependent firing profiles – all hallmarks of cell-intrinsic cold sensitivity that is independent of afferent synaptic input ^29, 30^. However, to date, the specific molecular and cellular identities of these cold-sensitive hypothalamic neurons and their role in mediating hypothermia-induced shivering thermogenesis remain unresolved.

In this study, we combined RNA-seq with projection-specific transcriptomic profiling to identify a distinct population of Calbindin D28k-expressing, VGLUT2-positive excitatory neurons in the DMH (DMH^Calb1/Vglut2^). CALB1 is a calcium-binding protein belonging to the calmodulin superfamily, which is essential for maintaining intracellular calcium homeostasis in neurons ^31, 32^. Using molecular genetics and electrophysiology, we demonstrate that DMH^Calb1/Vglut2^ neurons drive hypothermia-evoked overt shivering and exhibit intrinsic cold sensitivity, mediated by the transient A-type voltage-gated potassium currents (*I*_A_). Collectively, these data establish DMH^Calb1/Vglut2^ neurons as a dedicated, specific driver of hypothermia-induced shivering thermogenesis.

## Results

### Identification of DMH^Calb1/Vglut2^ neurons responsive to hypothermia

Prior behavioral and neuroprotection research demonstrated that ice-cold water exposure lowers mice’s T_core_ to hypothermia ^26, 33–35^. We adapted this approach by spraying ice-cold water onto the body surface of freely behaving mice to induce hypothermia and quantify associated shivering responses. At 25 °C ambient temperature (T_amb_), ice-cold water spray caused a sharp T_core_ drop, shivering rise, and stable hypothermia (Fig. 1A). Simultaneous T_core_ and posterior cervical muscle electromyography (EMG) showed a gradual EMG power (RMS) rise as T_core_ dropped 36.5–33°C.

**Fig. 1:**
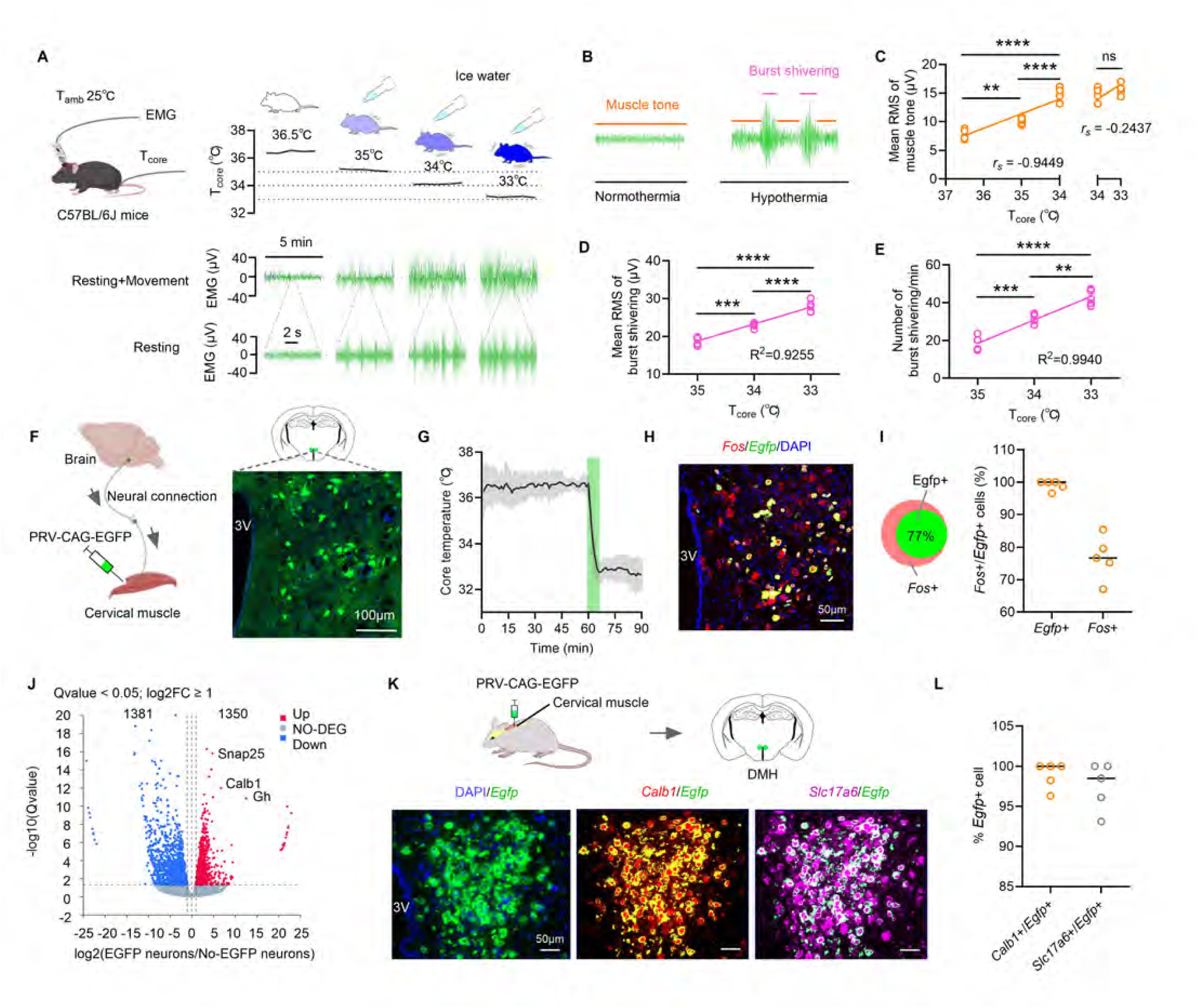
Combination of retrograde transsynaptic tracing and RNA-seq to identify DMH^Calb1/Vglut2^ neurons during hypothermia-induced shivering. (A) The schematic depicts simultaneous T_core_ and cervical intramuscular EMG recordings in freely moving mice held at 25 °C ambient temperature, with core temperature gradually lowered from 36.5 °C to 33 °C via ice-cold water spray cooling. (B) The cervical intramuscular EMG trace illustrates tonic muscle tone and phasic shivering bursts under normothermia and hypothermia in resting mice. (C) Spearman rank correlation between muscle tone, EMG power, and T_core_, with corresponding *r*_s_ values shown. (D) Pearson correlation coefficient of burst shivering EMG power versus T_core_ at 35 °C, 34 °C, and 33 °C, with corresponding R^2^ = 0.9255. (E) Pearson correlation coefficient of burst shivering counts versus T_core_ at 35 °C, 34 °C, and 33 °C, with corresponding R^2^ = 0.9940. (F) Schematic of multisynaptic retrograde tracing using PRV-CAG-EGFP, demonstrating anatomical connectivity between cervical muscles and the DMH. The right panel displays a micrograph of retrogradely labeled EGFP-positive neurons within the DMH; scale bar = 100 μm. (G) Mice received surface cooling via ice-cold water spray to reduce T_core_ to 33 °C for 30 min prior to brain harvesting for FISH analysis; n = 5 mice. (H) Representative FISH micrograph showing *Fos* mRNA induction in retrogradely traced *Egfp*⁺ DMH neurons after 30 min cooling to a T_core_ of 33 °C as described in (G); scale bar = 50 μm. (I) Quantification of the proportion of *Fos*⁺/*Egfp*⁺ double-positive neurons relative to total *Fos*⁺ or total *Egfp*⁺ neurons from images in (H). (J) Volcano plot from RNA-seq analysis displaying differentially expressed genes between EGFP⁺ and EGFP⁻ neurons (q < 0.05, Log₂ fold change ≥ 1); three biological replicates per group. (K) Schematics of retrograde tracing and representative FISH images from PRV-CAG-EGFP retrograde tracing of cervical muscles, demonstrating co-localization of *Calb1* and *SLC17A6* transcripts in EGFP⁺ DMH neurons on the same section; scale bars = 50 μm. (L) Quantification of the percentage of *Egfp*⁺ neurons co-expressing *Calb1* or *SLC17A6*, based on panel (K). Data were presented as means ± SD, with n = 5 mice. Correlation analyses were performed using Spearman’s (C) and Pearson’s correlation tests (D, E). One-way ANOVA with Tukey’s *post hoc* test was applied to panels C-E. ** p < 0.01, *** p < 0.001, **** p < 0.0001.

Resting state EMG was distinguished from movement and shivering-related rhythmic activity with synchronous video monitoring of behavioral movements (Extended Data Fig. 1A, B). Using MATLAB power analysis, we decomposed hypothermic EMG into tonic muscle tone and phasic shivering bursts (Extended Data Fig. 1A, B, and Fig. 1B). As T_core_ declined from 36.5 °C to 34 °C, muscle tone EMG power rose sharply (*r_s_* = −0.9449, p < 0.0001) then plateaued from 34 °C to 33 °C with no further change (*r_s_* = 0.2437, p = 0.5476; Fig. 1C), whereas tone frequency remained unchanged across cooling (Extended Data Fig. 1C–E). Phasic shivering only commenced when T_core_ dropped 35 °C below (hypothermia); its EMG power or shivering burst frequency increased linearly down to 33 °C (R^2^ = 0.9255 in Fig. 1D; R^2^ = 0.9940 in Fig. 1E). In summary, shivering is selectively triggered by hypothermia, while muscle tone serves as a steady baseline thermoregulatory mechanism.

To pinpoint hypothalamic nuclei governing shivering, we performed transsynaptic retrograde tracing from posterior cervical muscles to the brain with EGFP-expressing pseudorabies virus (PRV). Distinct DMH neurons exhibited strong EGFP labeling (Fig. 1F). We then lowered the mouse’s T_core_ to 33 °C for 30 min via ice-water spraying to test neuronal activation during hypothermia (Fig. 1G). Fluorescence *in situ* hybridization (FISH) confirmed that all PRV-EGFP-traced DMH neurons expressed *Fos* mRNA (Fig. 1H, I). Around 23% of unlabeled DMH cells were also *Fos*-positive, revealing that the DMH hosts a broader hypothermia-responsive neuronal pool, with muscle-projecting traced neurons constituting only a subpopulation (Fig. 1H, I).

To characterize PRV-EGFP-labeled DMH neurons, we sorted EGFP-positive neurons via fluorescence-activated cell sorting (FACS), using EGFP-negative cells as controls (Extended Data Fig. 2A, B). RNA-seq data revealed 1350 significantly upregulated genes in traced neurons (Fig. 1J, Extended Data Fig. 2C). Enrichment analysis highlighted excitatory neuron markers, notably *Slc17a6* (VGLUT2), alongside elevated glutamate transport and synaptic trafficking genes (*Slc1a3*, *Snap25*, *Stx1a*; Extended Data Fig. 2D). *Calb1* was also highly enriched in EGFP-positive cells (Fig. 1J, Extended Data Fig. 2E). Thus, DMH neurons innervating posterior cervical muscles are primarily VGLUT2/CALB1-expressing excitatory neurons.

We used FISH to characterize PRV-EGFP retrogradely-labeled DMH neurons. All EGFP-positive cells co-expressed *Slc17a6* and *Calb1* (Fig. 1K, L), yet no EGFP overlap was detected with inhibitory marker *Slc32a1* (VGAT; Extended Data Fig. 2F, G), verifying their excitatory identity. Immunofluorescence further confirmed EGFP-CALB1 colocalization (Extended Data Fig. 2H). In summary, traced DMH neurons are excitatory VGLUT2-expressing cells with high CALB1 levels, termed DMH^Calb1/Vglut2^.

### Hypothermia activation of DMH^Calb1/Vglut2^ neurons

We specifically labeled DMH^Calb1/Vglut2^ neurons in Calb1-IRES2-Cre-D (*Calb1^Cre^*) mice via stereotaxic injection of a recombinant adeno-associated virus vector (rAAV) carrying VGLUT2-DIO-EGFP elements into the DMH. Complete viral targeting was verified by full co-localization of *Egfp* with *Calb1* and *Slc17a6* transcripts (Extended Data Fig. 2I). These EGFP-tagged neurons permitted activity assessment during cold exposure. We also injected AAV2/9-VGLUT2-DIO-GCaMP6s into *Calb1^Cre^* mice for calcium fiber photometry recordings.

First, combining fiber photometry calcium recording and T_core_ tracking in freely moving mice (Fig. 2A), we found that T_core_ gradually dropped from 36.5 °C to 33 °C, GCaMP6s signals in DMH^Calb1/Vglut2^ neurons rose synchronously and incrementally (Fig. 2B, C). Correlation analysis showed a strong linear association between falling T_core_ and elevated calcium activity (R^2^ = 0.9092; Fig. 2C), demonstrating activation of DMH^Calb1/Vglut2^ neurons under hypothermic conditions.

**Fig. 2:**
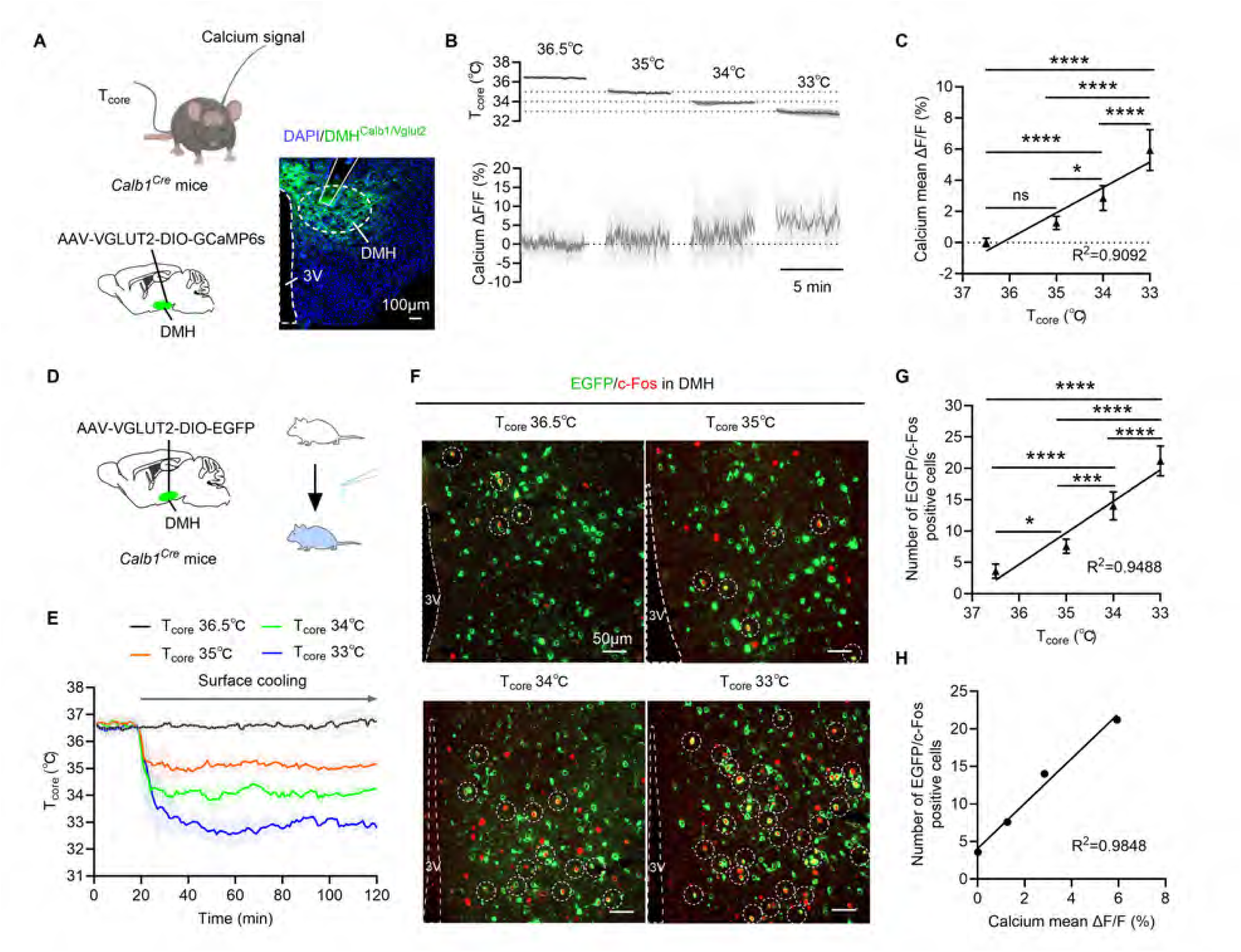
Hypothermia activates DMH^Calb1/Vglut2^ neurons. (A) Schematic illustrating simultaneous recordings of T_core_ and DMH calcium signals via fiber photometry in freely moving *Calb1^Cre^* mice. DMH^Calb1/Vglut2^ neurons were targeted with AAV-VGLUT2-DIO-GCaMP6s. A representative image is shown in the right-hand panel; n = 6 mice; scale bar = 50 μm. (B) Simultaneous recordings (setup in panel A) track T_core_ and calcium activity during ice-cold water spray-induced hypothermia from 36.5 °C down to 33 °C. (C) Quantification of changes in DMH^Calb1/Vglut2^ neuronal calcium signal (ΔF/F) and Pearson correlation analysis between T_core_ and ΔF/F values (data from panel B). (D) Schematic depicting stereotaxic delivery of AAV-VGLUT2-DIO-EGFP into the DMH of *Calb1^Cre^* mice to selectively label DMH^Calb1/Vglut2^ neurons with EGFP. (E) Mice prepared as described in panel D received ice-cold water spray to lower T_core_ to 35 °C, 34 °C, or 33 °C, with each temperature held for ∼90 min. (F) Representative immunofluorescence images showing c-Fos (red) expression colocalized with EGFP-tagged DMH^Calb1/Vglut2^ neurons in mice from experimental groups D and E; scale bars = 50 μm. (G) Quantification of EGFP⁺/c-Fos⁺ double-labeled neurons, together with Pearson correlation analysis correlating double-positive cell counts with T_core_. (H) Pearson correlation analysis examining the relationship between changes in calcium ΔF/F (%) and counts of EGFP⁺/c-Fos⁺ double-positive neurons (data from panels C and G). Data were presented as means ± SD. Correlation analyses were performed using Pearson correlation analysis for panels C, G, H. One-way ANOVA with Tukey’s *post hoc* test was used for panels C and G. * p < 0.05, ** p < 0.01, *** p < 0.001, **** p < 0.0001.

Second, we labeled DMH^Calb1/Vglut2^ neurons with EGFP in *Calb1^Cre^* mice via DMH-targeted AAV2/9-VGLUT2-DIO-EGFP injection (Fig. 2D). Cooling from 36.5°C to 33°C linearly increased c-Fos/EGFP co-labeled neurons (R² = 0.9488; Fig. 2E–G). The abundance of activated neurons also linearly correlated with GCaMP signal intensity during cooling (R² = 0.9848; Fig. 2H). At 33°C, about 81.7% of DMH c-Fos-positive neurons were DMH^Calb1/Vglut2^ cells (Fig. 2F). These data demonstrate that T_core_ reduction progressively activates DMH^Calb1/Vglut2^ neurons.

Lastly, DMH neurons are essential for shivering thermogenesis. Bilateral DMH electrolytic lesions lowered T_core_ relative to sham controls (Extended Data Fig. 3A–D) and suppressed muscle tone EMG power without altering its frequency (Extended Data Fig. 3E–G). Upon cooling to 33 °C, lesioned mice exhibited reduced muscle tone, burst-shivering EMG power, and burst-shivering frequency, while muscle tone frequency stayed unaltered (Extended Data Fig. 3H–J). Thus, the DMH modulates the magnitude of muscle tone and burst shivering, acting as a central hub governing shivering thermogenesis to maintain core temperature homeostasis.

### DMH^Calb1/Vglut2^ neurons specifically regulate hypothermia-evoked shivering

We applied Cre-dependent chemogenetics to dissect the specific function of DMH^Calb1/Vglut2^ neurons in hypothermia-induced shivering at T_amb_ of 25°C (Fig. 3A, 4A). For Cre-on manipulations, we injected into the DMH with the following rAAVs: AAV-VGLUT2-DIO-EGFP (control), AAV-VGLUT2-DIO-hM3D(Gq)-EGFP (activation), and AAV-VGLUT2-DIO-hM4D(Gi)-EGFP (inhibition) in *Calb1^Cre^* mice. The EGFP-positive neurons represent DMH^Calb1/Vglut2^ neurons (Fig. 3A). For Cre-off mice, *Calb1^Cre^* mice were injected in the DMH with the following rAAVs: AAV-VGLUT2-DO-mCherry (control), AAV-VGLUT2-DO-hM3D(Gq)-mCherry (activation), and AAV-VGLUT2-DO-hM4D(Gi)-mCherry (inhibition). The mCherry-positive neurons represent DMH^Calb1**-** /Vglut2+^ neurons (Fig. 4A). We then compared thermogenic outputs to verify selective shivering regulation by DMH^Calb1/Vglut2^ neurons.

**Fig. 3:**
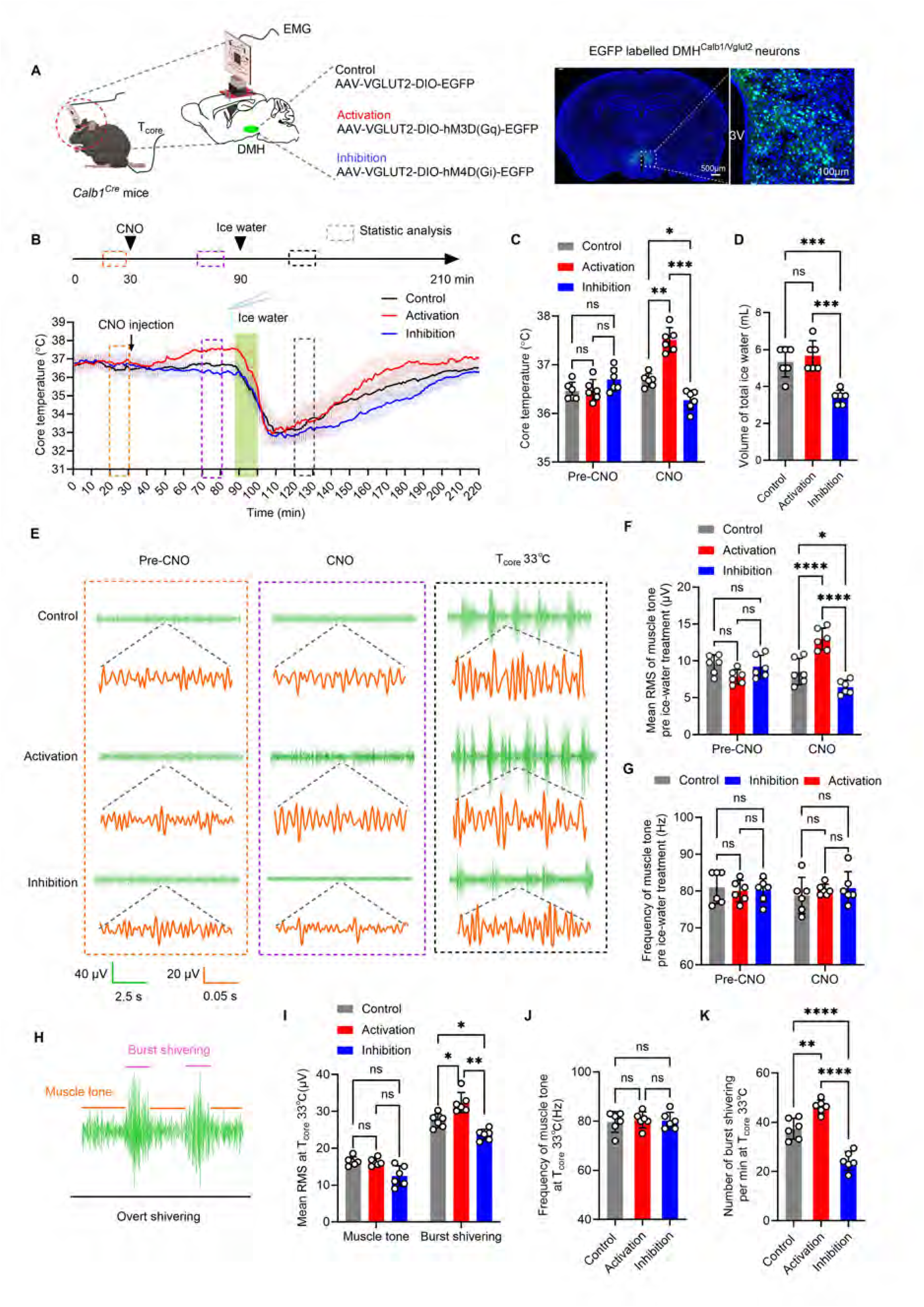
Chemogenetic modulation of DMH^Calb1/Vglut2^ neurons specifically affects hypothermia-evoked shivering. (A) Schematic depicting stereotaxic delivery of EGFP-labeled chemogenetic viruses into the DMH of *Calb1^Cre^* mice to specifically mark DMH^Calb1/Vglut2^ neurons. Simultaneous recordings of core temperature and EMG activity were conducted in freely moving animals. Coronal brain section micrographs on the right display EGFP fluorescence, confirming accurate viral injection targeting the bilateral DMH, with scale bars indicated on corresponding images. (B) Continuous T_core_ traces of mice injected with control virus (black line), chemogenetic activating virus (red line), and chemogenetic inhibitory virus (blue line). Vertical dashed boxes mark analysis time windows: pre-CNO phase (orange dashed line), CNO treatment phase (purple dashed line), and the time when T_core_ dropped to 33 °C (black dashed line). The horizontal green bar indicates the 10-minute duration of ice-cold water spray cooling. (C) Quantitative analysis of T_core_ variations during the 10-minute pre-CNO baseline period and the 10-minute CNO treatment period. (D) Quantification of the volume of ice-cold water spray needed to reduce T_core_ to 33 °C during the 10-minute cooling period indicated by the green bar in panel B. (E) Representative cervical intramuscular EMG traces acquired from a resting mouse during the time window specified in panel B. The green trace corresponds to a 10 s continuous recording, while the orange trace depicts a 300 ms segment of EMG activity reflecting basal muscle tone. The phasic burst shivering EMG signals are only visible at T_core_ of 33 °C (black dashed line). (F) Quantification of basal muscle tone EMG power measured pre-ice water treatment, analyzed across the time intervals defined in panel B. (G) Quantification of basal muscle tone EMG frequency (defined as the number of peaks of EMG signal per second) measured pre-ice water treatment, analyzed across the time intervals defined in panel B. (H) Representative cervical intramuscular EMG profile illustrating hypothermia-evoked overt burst shivering in resting mice, composed of basal muscle tone and phasic burst-type shivering activity. (I-K) Quantification of muscle tone and burst shivering EMG power (I), muscle tone EMG frequency (J), and burst shivering count (K) at T_core_ of 33 °C, 10 min following ice-cold water spray cooling per the time course in panel B. Data were expressed as mean ± SD (n = 6 mice). Two-way ANOVA followed by Tukey’s *post hoc* test was applied for analyses in panels C, F, and G; one-way ANOVA with Tukey’s post hoc test was used for panels D, I, J, and K. *p < 0.05, **p < 0.01, ***p < 0.001, ****p < 0.0001.

**Fig. 4:**
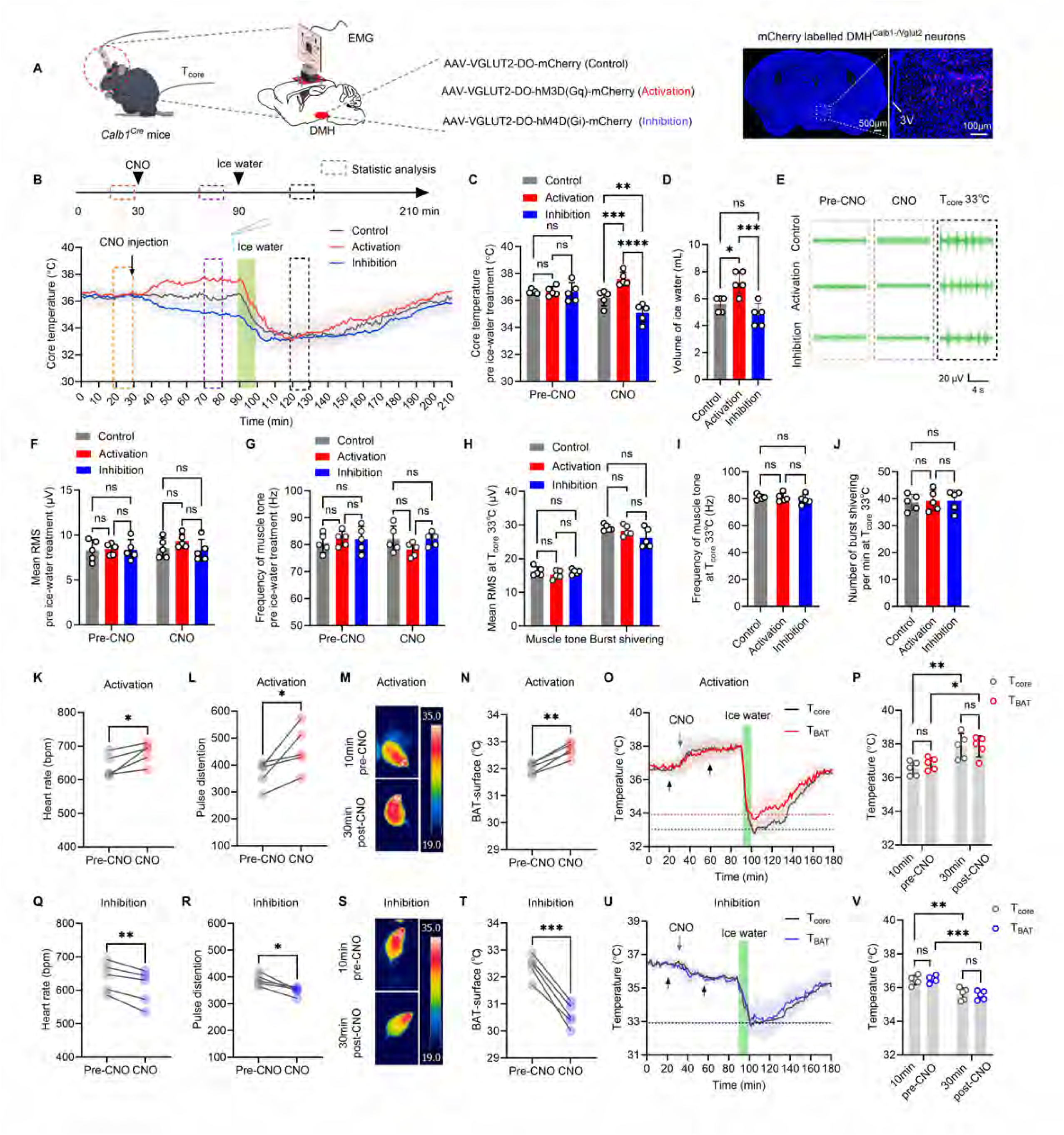
Chemogenetic modulation of Calb1- and Vglut2+ DMH neurons did not affect hypothermia-evoked shivering. (A) Schematic illustrating stereotaxic injection of mCherry-tagged chemogenetic viral constructs into the DMH of *Calb1^Cre^* mice to label DMH^Calb1⁻/Vglut2+^ neurons. Simultaneous T_core_ and EMG recordings were acquired in freely moving mice. Representative micrographs on the right confirm viral targeting within the DMH via mCherry red fluorescence. (B) Continuous recordings of T_core_ from mice injected with the control virus (black line), activation virus (red line), and inhibition virus (blue line). Dashed boxes indicate pre-CNO, CNO, and 33 °C T_core_ analysis windows; the green bar shows a 10-minute ice-cold water spray cooling period. (C) Quantified T_core_ shifts during the 10 min pre-CNO and 10 min CNO periods defined in (B). (D) Total ice-cold water spray volume needed to lower T_core_ to 33 °C within the 10 min cooling window in (B). (E) Representative cervical muscle EMG traces at baseline within the time frame in (B). (F) Baseline EMG power quantified across the intervals in (B) before ice-cold water spray. (G) Baseline EMG frequency quantified across the intervals in (B) before ice-cold water spray. (H-J) Quantified muscle tone EMG power and shivering burst EMG power (H), muscle tone EMG frequency (I), and shivering burst count (J) once T_core_ reached 33 °C at 10 min post ice-cold water spray, as indicated in (B). (K, L, Q, R) Heart rate (K, Q) and pulse distension (L, R) were quantified 10 min pre-CNO and 30 min post-CNO, following chemogenetic activation (K, L) or inhibition (Q, R) of DMH^Calb1⁻/Vglut2+^ neurons in free-moving mice. (M, N, S, T) Representative infrared interscapular BAT (iBAT) temperature (T_BAT_) images (M, S) and corresponding quantification (N, T), measured 10 min pre-and 30 min post-CNO during DMH^Calb1⁻/Vglut2+^ neuron activation or inhibition. (O, U) Simultaneous thermocouple recordings of T_core_ (black) and iBAT T_BAT_ (red/blue) in freely moving mice, before and after ice-cold water spray-induced cooling to 33 °C, under chemogenetic activation or inhibition of DMH^Calb1⁻/Vglut2+^ neurons. (P, V) Summary quantification of T_core_ and T_BAT_ at 10 min pre-CNO and 30 min post-CNO from traces (O) and (U), respectively, as indicated by the black arrows. Data were expressed as mean ± SD, with n = 5 mice per group. Two-way ANOVA followed by Tukey’s *post hoc* test was applied for panels C, F, G, P, and V. One-way ANOVA with Tukey’s *post hoc* correction was used for panels D, H, I, and J. Paired Student’s t-tests were performed for panels K, L, N, Q, R, and T. Significance levels: *p < 0.05, **p < 0.01, ***p < 0.001, ****p < 0.0001.

CNO treatment had no baseline effects on T_core_, heart rate, pulse distention, or T_BAT_ activity in control animals, as confirmed by infrared imaging, interscapular BAT thermocouple, and rectal temperature recordings (Extended Data Fig. 4).

Chemogenetic activation of Cre-on DMH^Calb1/Vglut2^ neurons raised T_core_ by 0.7 ± 0.06 °C (Fig. 3A-C) and increased muscle tone EMG power (without altering EMG frequency) (Fig. 3E-G and Extended Data Fig. 5A, B), while heart rate, pulse distention (Extended Data Fig. 5C, D), and T_BAT_ remained unchanged 30 min after CNO administration (Extended Data Fig. 5E-H). Conversely, activating Cre-off DMH^Calb1-/Vglut2+^ neurons failed to modulate muscle tone EMG yet significantly elevated T_core_, heart rate, pulse distention, and T_BAT_ compared to pre-CNO levels (Fig. 4B-P and Extended Data Fig. 4Q, R).

Chemogenetic suppression of Cre-on DMH^Calb1/Vglut2^ neurons, referred to as the ’inhibition’ group in Fig. 3A, lowered T_core_ by 0.3 ± 0.1 °C (Fig. 3B, C) and decreased muscle tone EMG power with no change in EMG frequency (Fig. 3E-G and Extended Data Fig. 5A, B), leaving heart rate, pulse distention (Extended Data Fig. 5I, J), and T_BAT_ unaltered (Extended Data Fig. 5K-N). By comparison, inhibiting Cre-off DMH^Calb1⁻/Vglut2⁺^ neurons did not affect muscle tone EMG but reduced T, heart rate, pulse distention, and T_BAT_ (Fig. 4B-J, Q-V). Together, these data confirm DMH^Calb1/Vglut2^ neurons selectively control skeletal muscle thermogenesis for T_core_ homeostasis.

Under hypothermic challenge, activating Cre-on DMH^Calb1/Vglut2^ neurons needed an equivalent amount of ice-cold water spray to lower T_core_ to 33 °C as controls (Fig. 3D), with baseline muscle tone EMG, heart rate and pulse distention unchanged when reaching 33 °C (Fig. 3H-J and Extended Data Fig. 6A). At 33 °C, this activation significantly increased burst shivering EMG power and counts (Fig. 3I, K). In contrast, activating Cre-off DMH^Calb1⁻/Vglut2⁺^ neurons required more ice-cold water spray to lower T_core_ to 33°C (Fig. 4D), markedly increased heart rate and pulse distention (Extended Data Fig. 6B), yet exerted no influence on muscle tone or burst shivering activity at 33 °C (Fig. 4H-J). Conversely, inhibiting Cre-on DMH^Calb1/Vglut2^ neurons required less ice-cold water spray to reduce T_core_ to 33°C compared to controls (Fig. 3D), without altering muscle tone EMG, or cardiovascular readouts (Fig. 3I, J; Extended Data Fig. 6A).

These inhibitory Cre-on mice displayed drastically reduced burst shivering EMG power and event counts (Fig. 3I, K). Suppressing Cre-off DMH^Calb1⁻/Vglut2⁺^ neurons lowered heart rate and pulse distention (Extended Data Fig. 6B) but produced no alterations in muscle tone or burst shivering activity (Fig. 4H-J). Across all manipulations, T_BAT_ stayed consistently above T_core_ during cold exposure (Extended Data Fig. 5G, H, M, N). Inhibition of Cre-off DMH^Calb1**-**/Vglut2+^ neurons fully abolished cold-triggered T_BAT_, driving parallel declines in T_BAT_ and T_core_ (Fig. 4U, V). Collectively, the Cre-dependent chemogenetics data demonstrated that DMH^Calb1/Vglut2^ neurons specifically regulate shivering thermogenesis without affecting cold-induced NST-related heart rate, pulse distention, or T_BAT_ activity.

We then applied optogenetic inhibition to examine how DMH^Calb1/Vglut2^ neurons control shivering thermogenesis (Fig. 5A). As shown in Fig. 5C, D, F and G, when T_core_ was lowered to 33 °C, optogenetic inhibition (laser-on) of DMH^Calb1/Vglut2^ neurons significantly suppressed burst shivering EMG power and event number, but did not affect muscle tone EMG power relative to the laser-off condition (Fig. 5F), confirming that DMH^Calb1/Vglut2^ neurons drive hypothermia-evoked burst-shivering. Interestingly, at a normothermic T_core_ of 36.5 °C (Fig. 5B), laser-on also reduced EMG power in muscle tone compared to the laser-off group (Fig. 5D, E).

**Fig. 5:**
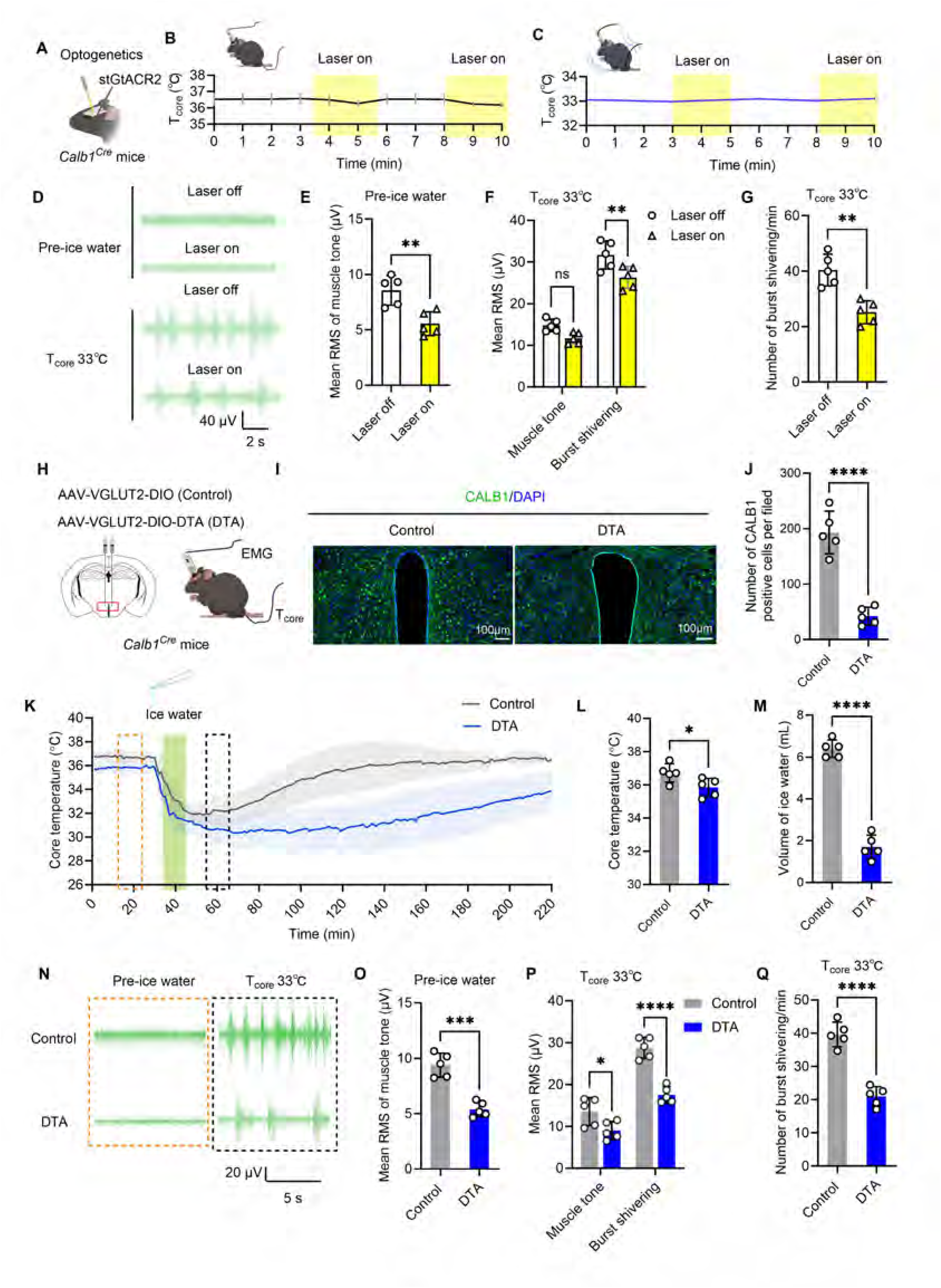
Inhibition of DMH^Calb1/Vglut2^ neurons alleviates hypothermia-induced shivering. (A-C) Three weeks following stereotaxic injection of optogenetic inhibitory virus (AAV2/9-VGLUT2-DIO-stGtACR2-EGFP-WPREs) into the DMH of *Calb1^Cre^* mice (A), T_core_ and posterior cervical muscle EMG were concurrently recorded during optogenetic suppression of DMH^Calb1/Vglut2^ neurons at baseline T_core_ of 36.5 °C (B) and hypothermic T_core_ of 33 °C (C); yellow and white intervals denote laser-on and laser-off epochs, respectively. (D) Representative cervical intramuscular EMG traces acquired during laser-on and laser-off periods defined in (B, C). (E) Quantification of muscle tone EMG power under laser-on and -off conditions prior to ice-cold water spray, as shown in panel D. (F, G) Quantitative comparison of muscle tone and shivering burst EMG power (F), as well as shivering burst counts (G), between laser-on and laser-off states at T_core_ of 33 °C. (H) Schematic depicting Cre-dependent DTA expression in the DMH of *Calb1^Cre^* mice for targeted ablation of DMH^Calb1/Vglut2^ neurons. (I) Representative CALB1 immunofluorescence staining counterstained with DAPI; scale bar = 100 μm. (J) Quantification of CALB1-positive neuronal density within the DMH from images in (I) (K) Continuous T_core_ recordings from freely moving DTA (blue) and control (black) mice, assessed at pre-cooling (orange dashed line) and once T_core_ decreased to 33 °C (black dashed line). (L) Quantitative analysis and statistical comparison of baseline T_core_ prior to surface cooling (orange dashed line in K). (M) Total ice-cold water spray volume required to lower T_core_ to 33 °C over the cooling window (green bar in K). (N) Representative resting cervical intramuscular EMG recordings across the experimental timeframe as indicated in (K). (O) Quantification of pre-ice-water-spray muscle tone EMG power derived from traces in (N). (P, Q) Quantification of muscle tone and shivering burst EMG power (P), and total shivering burst numbers (Q) at T_core_ of 33 °C. Data were presented as mean ± SD, with n = 5 mice. One-way ANOVA followed by Tukey’s *post hoc* test was used for panels F and P; paired t-tests were applied for panels E and G; unpaired t-tests were used for panels J, L, M, O, and Q. Statistical significance: *p < 0.05, **p < 0.01, ***p < 0.001, ****p < 0.0001.

Nevertheless, optogenetic inhibition (laser-on) in control virus-injected mice (AAV2/9-VGLUT2-DIO-EGFP-WPREs) produced no changes in posterior cervical muscle EMG power, burst shivering EMG power, or shivering counts at T_core_ of 36.5 °C or 33 °C (Extended Data Fig. 7) compared with the laser-off treatment, serving as a negative control. These results confirm that silencing DMH^Calb1/Vglut2^ neurons reduces hypothermia-induced shivering thermogenesis.

Collectively, DMH^Calb1/Vglut2^ neurons directly control shivering thermogenesis without affecting NST-related heart rate, pulse distention, or T_BAT_ activity. During hypothermia, DMH^Calb1/Vglut2^ neurons regulate the magnitude and frequency of burst-shivering events, while the muscle tone EMG power plateaus and remains unaffected. These findings establish DMH^Calb1/Vglut2^ neurons as a specialized group responsible for driving burst shivering.

### DMH^Calb1/Vglut2^ neurons are required for hypothermia-evoked shivering

We ablated DMH^Calb1/Vglut2^ neurons in *Calb1^Cre^* mice via Cre-dependent diphtheria toxin subunit A (DTA) expression (Fig. 5H), which depleted ∼80% of DMH CALB1-positive cells (Fig. 5I, J). Prior to the cold challenge, DTA mice had lower T_core_ and muscle tone EMG power than controls (Fig. 5K, L, N, O; Extended Data Fig. 8A, B). During cooling, these mice reached a core temperature of 33 °C with less ice-cold water spray, along with attenuated muscle tone, weaker burst shivering EMG activity, and fewer shivering events (Fig. 5P, Q; Extended Data Fig. 8A, B). These data reinforce that DMH^Calb1/Vglut2^ neurons regulate hypothermia-evoked shivering thermogenesis. Interestingly, their ablation had no effect on T_BAT_, which remained above T_core_ during hypothermia in DTA mice (Extended Data Fig. 8C), confirming that these neurons did not mediate cold-stimulated T_BAT_.

### Hypothermic shivering driven by DMH^Calb1/Vglut2^ neurons is independent of synaptic input

To determine whether hypothermia directly activates DMH^Calb1/Vglut2^ neurons to drive shivering, we blocked upstream thermoregulatory inputs via intra-DMH infusion of synaptic blockers or viral deletion of afferent neurons. As shown in Fig. 6A, 25 min after bilateral DMH delivery of synaptic transmission inhibitors containing DNQX, D-AP5, and gabazine, T_core_ increased by approximately 2 °C, accompanied by heightened muscle tone EMG power but unchanged EMG frequency compared to vehicle controls (Fig. 6B-F). No obvious burst shivering occurred before or after this treatment.

**Fig. 6.**
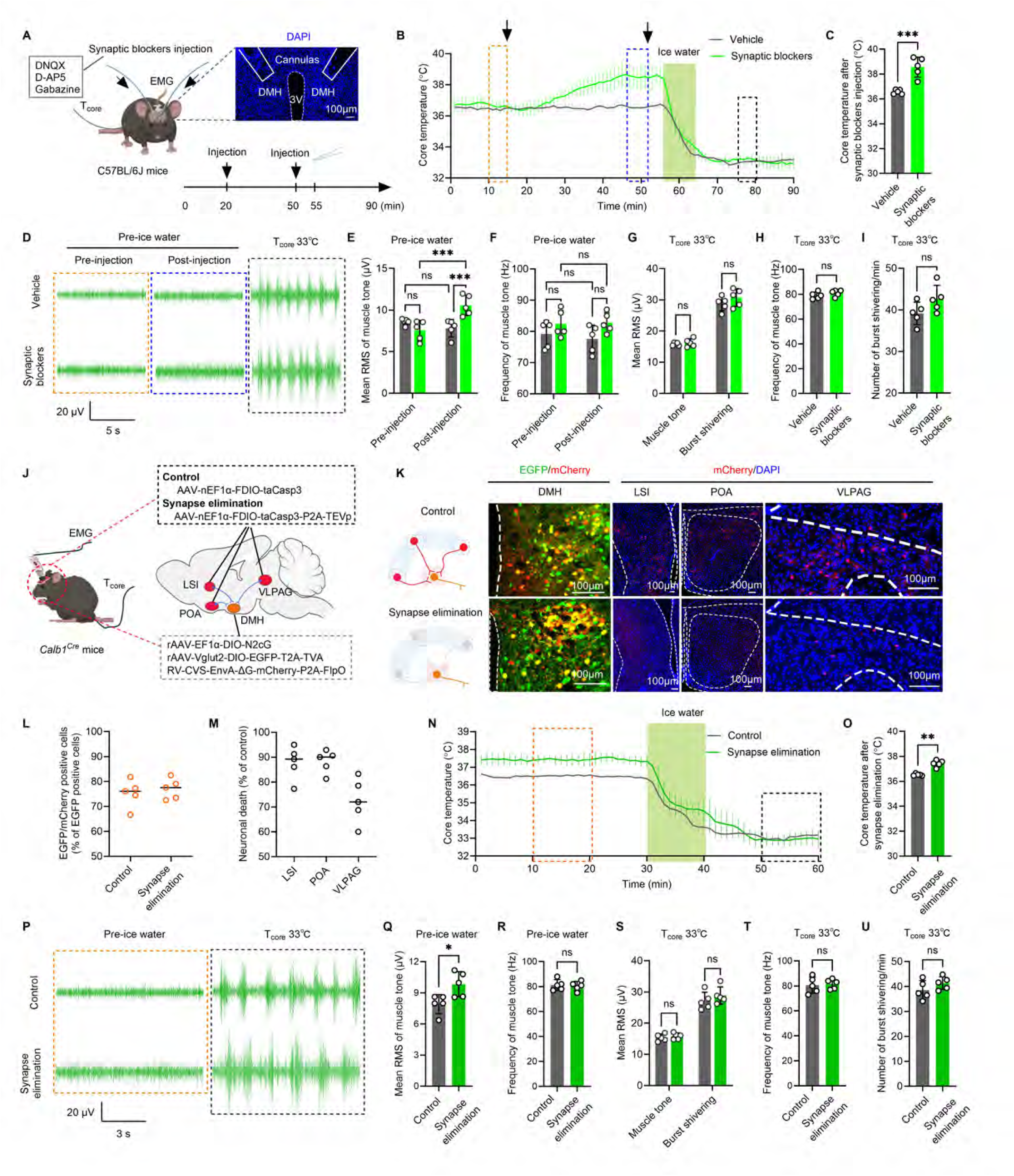
DMH^Calb1/Vglut2^ neuron activation of hypothermic shivering independent of synaptic input. (A) Schematic for T_core_ and EMG recordings before and after bilateral intra-DMH infusion of synaptic blockers (DNQX, D-AP5, gabazine) via implanted cannulae. DAPI-stained brain slices verified cannula injection placement. The experimental timeline is as illustrated. Scale bar = 100 μm. (B) Continuous recordings of T_core_ for freely moving mice receiving vehicle (black) or synaptic blocker infusion (green). Dashed boxes denote analysis epochs: pre-infusion (orange), post-infusion (blue), and T_core_ stabilized at 33 °C (black). Black arrows mark drug-delivery time points; the green bar represents a 10 min ice-cold water spray cooling protocol. (C) Quantification of T_core_ changes during the post-infusion window (blue dashed box in B). (D) Representative resting cervical intramuscular EMG recordings corresponding to the time windows in B. (E, F) Quantification of muscle tone EMG power (E) and frequency (F) prior to ice-cold water spray, depicted in D. (G-I) Quantification of muscle tone and shivering burst EMG power (G), muscle tone EMG frequency (H), and shivering burst counts (I) at T_core_ of 33 °C (J) Schematic of simultaneous T_core_ and EMG recordings in freely moving mice following viral-mediated ablation of presynaptic upstream neurons innervating DMH^Calb1/Vglut2^ neurons (K) Representative images from retrograde monosynaptic tracing targeting DMH^Calb1/Vglut2^ neurons (EGFP/mCherry double-positive). Control animals exhibit abundant mCherry-labeled upstream neurons in the LSI, POA and VLPAG; the synaptic ablation group shows marked loss of these mCherry-positive cells, consistent with caspase-3-driven apoptotic elimination. Scale bar = 100 μm. (L) Quantification of the fraction of EGFP⁺ DMH neurons co-expressing mCherry (from images in K). (M) The death rate of DMH-projecting upstream neurons was calculated as 100% minus neuronal survival percentage, where survival was defined as the ratio of mCherry⁺ neuron counts in the ablation group relative to controls. (N) Continuous recordings of T_core_ for mice with ablated upstream afferents to DMH^Calb1/Vglut2^ neurons (green) and control littermates (black). Dashed boxes indicate pre-cooling (orange) and T_core_ at 33 °C (black) analysis windows; the green bar denotes the 10 min ice-cold water spray cooling period. (O) Quantification of T_core_ within the pre-cooling period (orange dashed box in N). (P) Representative resting cervical intramuscular EMG traces matching the time intervals shown in panel N. (Q, R) Quantification of pre-cooling muscle tone EMG power (Q) and frequency (R) from traces in P. (S-U) Quantification of muscle tone and shivering burst EMG power (S), muscle tone EMG frequency (T), and total shivering burst numbers (U) at T_core_ of 33 °C. Data are presented as mean ± SD, n = 5 mice per group. Two-way ANOVA with Tukey’s *post-hoc* test was applied for panels E and F; one-way ANOVA with Tukey’s *post hoc* test for panels G and S; unpaired t-tests for panels C, H, I, O, Q, R, T and U. Significance: *p < 0.05, **p < 0.01, ***p < 0.001.

Subsequent cooling to hypothermic 33 °C with ice-cold water spray still triggered shivering 5 min after a second injection (Fig. 6D). However, there were no differences in the EMG powers of muscle tone, or burst shivering versus vehicle control (Fig. 6G-I). Overall, these findings demonstrate that hypothermia activates DMH^Calb1/Vglut2^ neurons and triggers shivering independently of upstream neural circuits.

We used retrograde monosynaptic rabies virus (RV) tracing in *Calb1^Cre^* mice via intra-DMH injection to identify upstream DMH^Calb1/Vglut2^ innervating neurons (Extended Data Fig. 9A). RV trans-synaptically labeled presynaptic neurons with EGFP- and tdTomato co-expression (Extended Data Fig. 9B, left panel). Whole-brain serial section imaging detected labeled inputs from the LSI, POA, and VLPAG (Extended Data Fig. 9B, right panel). FISH confirmed inhibitory neurons made up 94%, 88%, and 87% of these labeled populations in the LSI, POA, and VLPAG, respectively (Extended Data Fig. 9C-F).

To selectively ablate presynaptic inputs to DMH^Calb1/Vglut2^ neurons, we injected FDIO-taCasp3-TEVp virus into the POA, LSI, and VLPAG, followed by intra-DMH delivery of FlpO-expressing RV. Retrograde FlpO expression induced FDIO-dependent taCasp3/TEVp production in upstream neurons, triggering caspase-3 activation and targeted ablation of DMH-projecting cells; control virus lacked the TEVp caspase-activating cassette (Fig. 6J). DMH^Calb1/Vglut2^ neurons were labeled with EGFP. The fraction of EGFP⁺ neurons co-expressing mCherry was comparable between control (∼75%) and ablation (∼77%) groups (Fig. 6K, L). These double-labeled neurons permitted transsynaptic viral trafficking to upstream populations, driving mCherry expression in controls or caspase-dependent apoptosis in the ablation group (Fig. 6K). Quantification revealed ablation efficiencies of 87% in LSI, 88% in POA, and 73% in VLPAG relative to controls (Fig. 6K, M).

Ablating upstream inputs to DMH^Calb1/Vglut2^ neurons increased T_core_ and muscle tone EMG power without altering EMG frequency or triggering spontaneous shivering (Fig. 6N-R). Upon 33 °C hypothermia, shivering emerged equally in both the control and synapse-elimination groups (Fig. 6P), with no intergroup differences in quantified EMG parameters (Fig. 6S-U).

Taken together, these results showed that DMH^Calb1/Vglut2^ neurons receive mainly inhibitory presynaptic inputs. Ablation of these inputs increased muscle tone but did not produce overt shivering. Hypothermia at 33 °C still induced shivering regardless of upstream input suppression, demonstrating hypothermia directly activates DMH^Calb1/Vglut2^ neurons to initiate shivering, independent of upstream inhibitory disinhibition.

### DMH^Calb1/Vglut2^ neurons are hypothermia-sensitive

To determine direct cold sensing by DMH^Calb1/Vglut2^ neurons, we labeled them via AAV2/9-VGLUT2-DIO-EGFP injection into *Calb1^Cre^*mice. We performed whole-cell patch-clamp recordings on EGFP-positive DMH^Calb1/Vglut2^ neurons in the presence of synaptic blockers (DNQX, D-AP5, gabazine) to eliminate synaptic inputs. Using a precise temperature-controlled recording chamber, as shown in Extended Data Fig. 10A, the neuronal activity was recorded at 33 °C (cold), 36 °C, and 39 °C (warm) (Fig. 7A–C; Extended Data Fig. 10B).

**Fig. 7:**
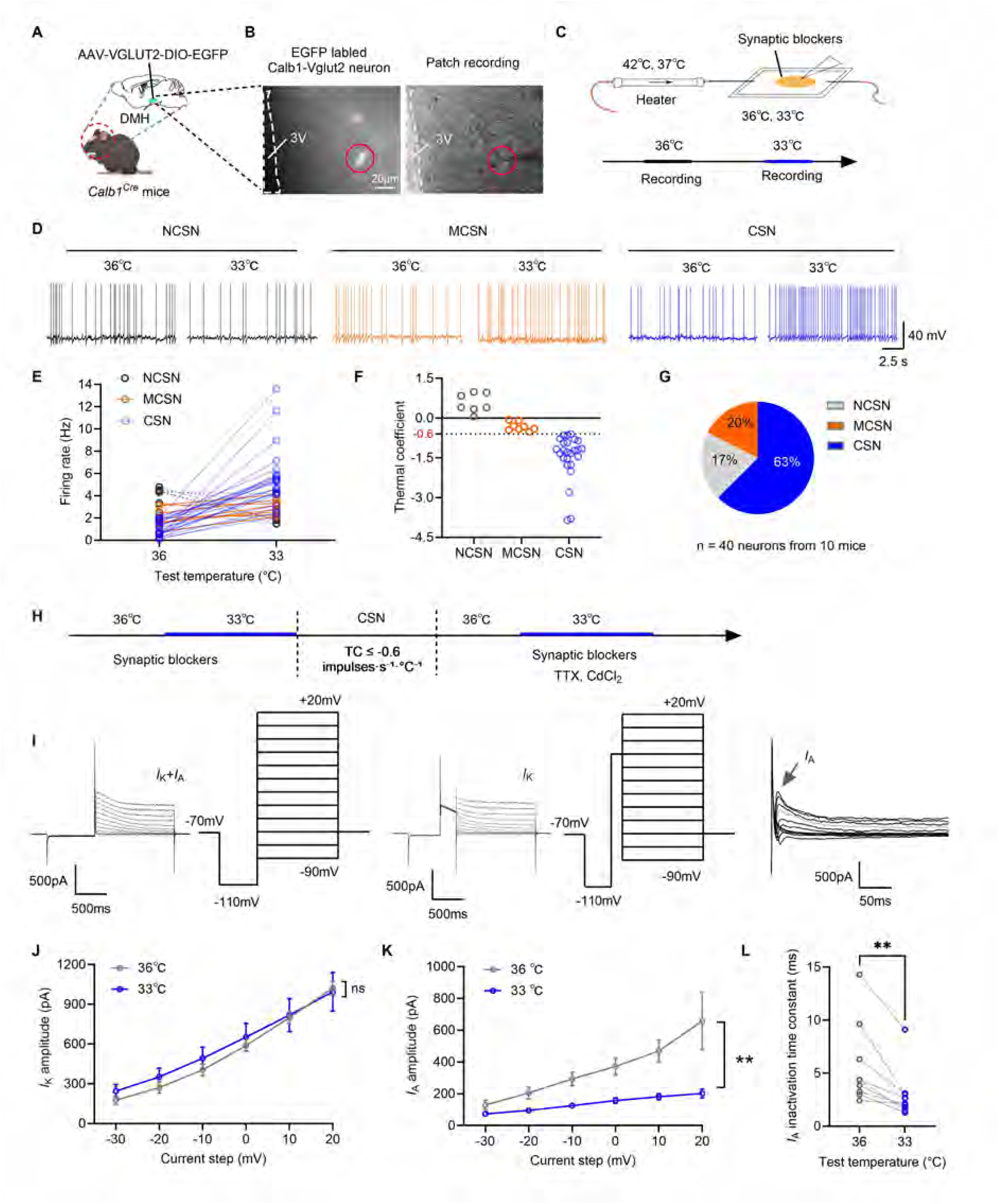
DMH^Calb1/Vglut2^ neurons are hypothermia-sensitive. (A) Schematic of stereotaxic injection of AAV-VGLUT2-DIO-EGFP into the DMH of *Calb1^Cre^* mice for specific EGFP expression in DMH^Calb1/Vglut2^ neurons. (B) Fluorescent micrograph of acute brain slices, showing EGFP-labeled DMH neurons (red circles) targeted for patch-clamp recordings. Scale bar = 20 μm. (C) Schematic of the patch-clamp recording setup with precise temperature control. The recording chamber was continuously perfused with aCSF containing synaptic blockers (20 μM DNQX, 50 μM D-AP5, and 20 μM gabazine). (D) Representative action potential traces of three subtypes of DMH neurons, including non-cold-sensitive neurons (NCSNs), moderately cold-sensitive neurons (MCSNs), and cold-sensitive neurons (CSNs), recorded at 36 °C and 33 °C. (E) Quantitative analysis of firing rates of NCSNs, MCSNs, and CSNs at 36 °C and 33 °C, corresponding to the traces in (D). (F) Temperature coefficient (TC) was calculated to quantify temperature-dependent changes in neuronal firing rate (impulses·s·°C). Neurons were classified based on TC values: NCSNs (TC > 0), MCSNs (0 > TC > −0.6), and CSNs (TC ≤ −0.6). (G) Proportions of NCSNs, MCSNs, and CSNs among total recorded DMH neurons (n = 40 neurons from 10 mice). (H) Schematic of the patch-clamp recording protocol. Recordings were initially performed at 36 °C and 33 °C in aCSF supplemented with synaptic blockers. After identification of CSNs (TC ≤ −0.6), recordings were repeated at both temperatures following additional perfusion with voltage-gated Na^+^ and Ca^2+^ channel blockers (1 μM TTX and 50 μM CdCl_2_). (I) Representative traces and recording protocols for sustained K^+^ current (*I*_K_) and transient K^+^ current (*I*_A_). (J) Quantitative analysis of sustained *I*_K_ amplitudes. (K) Quantitative analysis of transient *I*_A_ amplitudes. (L) Quantitative analysis of transient *I*_A_ inactivation time constant. Data were presented as mean ± SEM (J, K, L with n = 9 neurons from 4 mice). Repeated-measures two-way ANOVA with Tukey’s *post hoc* test was applied for statistical analysis in (J) and (K). Paired t-tests were applied for panel L. Significance: **p < 0.01.

Neuronal thermal sensitivity is defined by the thermal coefficient (TC), representing temperature-induced changes in firing rate (dF/dT = impulses·s⁻¹·°C⁻¹). A minimum TC of -0.6 was used to identify cold-sensitive neurons (CSN), with values between -0.6 and 0 indicating moderately cold-sensitive neurons (MCSN), and non-cold sensitive neurons (NCSN, TC > 0) per established criteria ^26, 36, 37^. Of 40 DMH^Calb1/Vglut2^ neurons recorded from 10 mice, 63% exhibited elevated firing as CSNs, 20% were MCSNs, and 17% were NCSNs (Fig. 7D-G). Additional recordings of 21 neurons across all subtypes showed no warm-sensitive responses at 39 °C (dF/dT > 0.8 Hz/°C; Extended Data Fig. 10C-H). Collectively, the majority of DMH^Calb1/Vglut2^ neurons are intrinsically cold-sensitive at hypothermic 33 °C.

To validate the specificity of this cold response, we recorded firing activity at 36 °C and 33 °C in neurons from the perifornical nucleus adjacent to the DMH as negative controls (Extended Data Fig. 11A, B). Perifornical neurons failed to display a temperature-dependent rise in firing rate and did not satisfy the TC criterion characteristic of cold-sensitive neurons. The TC values were all at dF/dT > 0 Hz/°C (n = 18 neurons; Extended Data Fig. 11C–E). This confirms that temperature-specific firing modulation is unique to the cold-responsive DMH^Calb1/Vglut2^ neurons.

### Voltage-gated K^+^ channels modulate DMH^Calb1/Vglut2^ neurons’ hypothermia sensitivity

To elucidate how hypothermia activates DMH^Calb1/Vglut2^ neurons, we reexamined RNA sequencing data of retrogradely labeled DMH neurons, which revealed differential expression of multiple potassium channel genes, including *Kcnab2*, *Kcne1l*, *Kcnh6*, and *Kcnk1* between EGFP-positive and EGFP-negative populations (Fig. 1F, J, and Extended Data Fig. 12A-C). Following this lead, we analyzed the properties of action potentials. Cooling to 33 °C did not alter spike threshold, amplitude, rise/decay kinetics, or afterhyperpolarization (AHP) duration (Extended Data Fig. 12D-J). However, 33 °C significantly broadened action potential half-width and reduced AHP amplitude relative to 36 °C (Extended Data Fig. 12K, L), two parameters closely associated with K⁺ efflux and neuronal hyperpolarization ^38, 39^.

Neuronal outward K⁺ currents consist of sustained delayed rectifier (*I*_K_) and transient A-type (*I*_A_) components. To examine their temperature-dependent modulation, during a patch-clamping experiment, we first identified CSNs based on the calculated TC values of DMH^Calb1/Vglut2^ neurons for their firing rates at 36 °C and 33 °C (Fig. 7H). Then the recorded *I*_K_ and *I*_A_ were measured using the voltage protocol illustrated in Fig. 7I, with Na⁺ and Ca²⁺ channels pharmacologically blocked by TTX and CdCl₂, respectively. Cooling to 33 °C significantly suppressed *I*_A_ but exerted no effect on *I*_K_ (Fig. 7J, K). Moreover, the *I*_A_ inactivation time constant was markedly shorter at 33 °C (hypothermia) relative to 36 °C. Collectively, these data indicate that hypothermic suppression of *I*_A_ magnitude, together with accelerated *I*_A_ inactivation, promotes membrane repolarization and enhances action potential firing in DMH^Calb1/Vglut2^ neurons, revealing a potential biophysical mechanism underlying their intrinsic cold sensitivity.

## Discussion

We identified DMH^Calb1/Vglut2^ neurons as selective mediators of hypothermia-triggered shivering thermogenesis in mice. These neurons function independently of canonical cutaneous cold-sensing and homeostatic thermoregulatory circuits, firing exclusively under hypothermic conditions with synaptic input-independent thermal thresholds. Mechanistically, hypothermia suppresses voltage-gated A-type K⁺ currents in these neurons, supporting their essential role in driving hypothermia-induced shivering thermogenesis.

Canonical thermoregulation theory holds that the brain lacks intrinsically cold-sensitive neurons, with body temperature homeostasis exclusively governed by POA warm-sensitive neurons (WSNs) first identified in the 1960s ^1, 40–42^. Cold-induced physiological responses are traditionally attributed to suppressed POA WSN activity or peripheral thermoreceptor signaling, rather than central cold-sensing neuronal activation. Despite advances in modern neurophysiological techniques, this long-standing dogma has remained unrefuted, as intact *in vivo* recordings cannot dissociate central neuronal cold sensitivity from confounding synaptic inputs of peripheral and central thermosensory pathways ^4, 42^.

This study identifies a group of intrinsically hypothermia-sensitive neurons within DMH. *Ex vivo* hypothalamic recordings with complete synaptic blockade confirmed that these neurons exhibit robust firing elevation under hypothermic conditions (< 35 °C), independent of peripheral thermoreceptor signals and upstream central synaptic inputs. Unlike canonical POA warm-sensitive neurons that sustain normothermic homeostasis by detecting minor core temperature fluctuations, DMH cold-sensitive neurons are selectively tuned to hypothermic thresholds and specifically mediate hypothermia-evoked shivering thermogenesis. These findings reveal an unrecognized parallel DMH thermosensory pathway that cooperates with classical thermoregulatory circuits to drive context-dependent thermal defense responses.

During hypothermia, low-amplitude muscle tone fills intervals between large shivering bursts on EMG (Extended Data Fig. 1B). Baseline resting muscle tone and overt shivering represent two distinct muscle thermogenic pathways for core temperature homeostasis. At ambient temperature, resting muscle tone generates heat via subtle subthreshold contractions. Chemogenetic inhibition/ablation of DMH^Calb1/Vglut2^ neurons reduced muscle tone power, whereas blocking upstream synaptic inputs increased muscle tone (Fig. 6), demonstrating baseline muscle tone is constitutively suppressed by inhibitory inputs within canonical thermoregulatory circuits. Only severe cold (<7 °C) triggers robust shivering to rapidly generate heat ^15, 17^.

Since muscle tone also mediates posture, arousal, and nociception, its thermogenic contribution during hypothermia remains understudied ^15, 17, 18^. Plateaus in muscle tone frequency and amplitude during cooling (Extended Data Fig. 1C-E and Fig. 1C) imply that tone-derived heat accounts for only a small fraction of total cold-induced thermogenesis. Future work should characterize additional DMH populations that regulate muscle tone and their crosstalk with DMH^Calb1/Vglut2^ neurons for the maintenance of normothermia. The body deploys distinct cold-defense mechanisms at different temperature thresholds, reflecting a hierarchical, energy-dependent response _43, 44._

Thermosensitive ion channels and receptors convert ambient thermal stimuli into neuronal electrical activity, constituting the core conserved molecular mechanism for cellular temperature detection across nervous systems ^45^. Distinct thermally responsive ion channels and receptors possess divergent activation temperature thresholds, enabling neurons to discriminate graded thermal signals ranging from mild warmth to severe cold. Among thermoregulatory ion channel families, voltage-gated potassium channels occupy a critical functional position in shaping neuronal thermal responsiveness ^46^. Notably, the KCNQ1 subtype exhibits pronounced sensitivity to mild cold within the physiological range of 25–35LJ°C, a temperature range overlapping with thermal fluctuations that trigger thermogenesis in mice ^47, 48^. However, KCNQ1-dependent mild cold responses are unlikely to drive phasic shivering; instead, they may affect elevated muscle tone, although this putative pathway remains unconfirmed.

Our electrophysiological recordings uncover a distinct biophysical mechanism underlying the intrinsic cold sensitivity of DMH^Calb1/Vglut2^ neurons. Unlike canonical warm-sensitive neurons in the preoptic area and hypothalamus, which exhibit suppressed excitability under cooling, these DMH neurons are selectively modulated by atypical temperature-dependent inhibition of the transient outward potassium current *I*_A_, with the delayed rectifier potassium current *I*_K_ functionally unaffected. Cooling from 35 °C to 33 °C drives the activation of DMH^Calb1/Vglut2^ neurons primarily through *I*_A_-mediated remodeling of action potential kinetics. Shortened action potential decay and AHP duration synergistically boost cold-induced neuronal excitability ^49^.

Transcriptomic evidence corroborates this functional phenotype by showing elevated expression of potassium channel isoforms in DMH^Calb1/Vglut2^ neurons compared with other DMH subtypes. Based on these findings, we propose a plausible mechanism by which hypothermia relieves the *I*_A_-mediated inhibitory brake by dampening *I*_A_ amplitude and accelerating its inactivation. Concurrent intact *I*_K_ maintains efficient spike repolarization, thereby curtailing inter-spike intervals and sustaining high-frequency firing during hypothermia.

This study explored shivering neural mechanisms using freely behaving awake mice, whose EMG recordings are susceptible to motion artifacts. Three approaches were applied to acquire and analyze resting-state EMG signals: (1) Habituation: Mice underwent 2 h daily acclimation for two consecutive days before testing, plus a 30-min pre-test adaptation. All trials were performed in quiet conditions by the same researchers between 13:00 and 17:00 to stabilize the animals for precise EMG measurement. (2) Baseline differentiation: Limb movement elevates EMG amplitude, which can be clearly distinguished from resting baseline signals. (3) Cold response separation: Ice-cold water-induced hypothermia triggers shivering, whereas locomotion terminates shivering and generates distinct motion-specific EMG waveforms, enabling independent quantification of shivering-related signals. This workflow enables reliable resting-state EMG acquisition in freely moving mice.

Decoding central shivering regulation holds critical biological and clinical value ^26, 50–52^. For example, therapeutic hypothermia at 33 °C protects against brain injuries yet is hampered by strong shivering, which delays cooling and worsens organ damage ^26, 51, 53–55^. Current anti-shivering drugs carry risks like respiratory depression and hypotension ^56–58^. Targeting DMH^Calb1/Vglut2^ neurons to suppress shivering without cardiovascular side effects offers a safer strategy for optimized hypothermic therapy and warrants further study.

## Limitations of the study

This study focused on identifying shivering-regulating neuronal subpopulations within the DMH. The function of cold-insensitive DMH^Calb1/Vglut2^ neurons in regulating muscle tone is unconfirmed. Moreover, it is unclear whether the three thermally distinct neuron subsets correspond to potassium channel-linked functional subtypes with separate temperature-sensing modules. Future work should clarify the roles of cold-insensitive neurons and the molecular mechanisms driving differential muscle shivering responses to cold.

## Online content

Any methods, additional references, Nature Portfolio reporting summaries, source data, extended data, supplementary information, acknowledgments, peer review information, details of author contributions and competing interests, and statements of data and code availability are available at XXXX online.

## Methods

### Key resources table

| REAGENT or RESOURCE | SOURCE | IDENTIFIER |
| --- | --- | --- |
| **Antibodies** |  |  |
| Mouse anti-c-Fos | Abcam | Cat#ab208942; RRID: AB_2747772 |
| Rabbit anti-c-Fos | Synaptic Systems | Cat#226008; RRID: AB_2891278 |
| Rabbit anti-Calbindin-D28k | Proteintech | Cat#14479-1-AP; RRID: AB_2228318 |
| 594 goat anti-mouse | Abcam | Cat#ab150116; RRID: AB_2650601 |
| 594 goat anti-rabbit | Abcam | Cat#ab150080; RRID: AB_2650602 |
| 488 goat anti-rabbit | Abcam | Cat#ab150077; RRID: AB_2630356 |
| **Bacterial and virus strains** |  |  |
| PRV-CAG-EGFP | BrainVTA, Wuhan | Cat#P03001 |
| AAV2/9-VGLUT2-DIO-hM3D(Gq)-EGFP-WPREs | BrainVTA, Wuhan | Cat#PT-4801 |
| AAV2/9-VGLUT2-DIO-hM4D(Gi)-EGFP-WPREs | BrainVTA, Wuhan | Cat#PT-3882 |
| AAV2/9-VGLUT2-DO-hM3D(Gq)-mCherry-WPREs | BrainVTA, Wuhan | Cat#PT-13917 |
| AAV2/9-VGLUT2-DO-hM4D(Gi)-mCherry-WPREs | BrainVTA, Wuhan | Cat#PT-13916 |
| AAV2/9-VGLUT2-DIO-DTA-mCherry-WPRE-hGH polyA | BrainVTA, Wuhan | Cat#PT-11519 |
| AAV2/9-VGLUT2-DIO-Gcamp6s-WPREs | BrainVTA, Wuhan | Cat#PT-8348 |
| AAV2/9-VGLUT2-DIO-stGtACR2-EGFP-WPREs | BrainVTA, Wuhan | Cat#PT-13468 |
| AAV2/9-VGLUT2-DIO-EGFP-WPREs | BrainVTA, Wuhan | Cat#PT-3884 |
| AAV2/9-VGLUT2-DO-mCherry-WPREs | BrainVTA, Wuhan | Cat#PT-13918 |
| rAAV-EF1α-DIO-N2cG | Brain Case | Cat#BC-0042-9 |
| rAAV-VGLUT2-DIO-EGFP-T2A-TVA | Brain Case | Cat#BC-3580 |
| RV-CVS-EnvA-ΔG-tdTomato | Brain Case | Cat#BC-RV-CVS-EnvA462 |
| AAV-nEF1α-FDIO-taCasp3-P2A-TEVp | Brain Case | Cat#BC-2001 |
| AAV-nEF1α-FDIO-taCasp3 | Brain Case | Cat#BC-2002 |
| RV-CVS-EnvA-ΔG-mCherry-P2A-FlpO | Brain Case | Cat#BC-RV-CVS-EnvA471 |
| **Chemicals, peptides, and recombinant proteins** |  |  |
| Meloxicam | MedChemExpress LLC | Cat#HY-B0261; CAS: 71125-38-7 |
| Clozapine N-oxide | MedChemExpress LLC | Cat#HY-17366; CAS: 34233-69-7 |
| DNQX | MedChemExpress LLC | Cat#HY-15067; CAS: 2379-57-9 |
| Gabazine | MedChemExpress LLC | Cat#HY-103533; CAS: 104104-50-9 |
| D-AP5 | MedChemExpress LLC | Cat#HY-100714A; CAS: 79055-68-8 |
| Cadmium chloride (CdCl<sub>2</sub>) | Sigma-Aldrich | Cat#202908; CAS: 10108-64-2 |
| Tetrodotoxin (TTX) | MEIZHENG BIO-TECH | Cat#MSS0002; CAS: 4368-28-9 |
| Isoflurane | RWD Life Science | Cat#R510-22-10 |
| Crystal violet | Sigma-Aldrich | Cat#C5042; CAS: 10510-54-0 |
| Critical commercial assays |  |  |
| RNAscope Multiplex Fluorescent Assays V2 | Advanced Cell Diagnostics | 323110 |
| Adult Brain Neuron Isolation Kit | Miltenyi Biotec | 130-107-677 |
| Experimental models: Organisms/strains |  |  |
| Mouse: Calb1-cre | Jackson Laboratory | RRID: IMSR_JAX:028532 |
| Mouse: C57BL/6J | Vital River Laboratory Animal Technology Co., Ltd. | N/A |
| Oligonucleotides |  |  |
| Slc17a6 in situ hybridization probe | Advanced Cell Diagnostics | 1170921-C1 |
| Egfp in situ hybridization probe | Advanced Cell Diagnostics | 538851-C2 |
| tdTomato in situ hybridization probe | Advanced Cell Diagnostics | 317041-C2 |
| Calb1 in situ hybridization probe | Advanced Cell Diagnostics | 428431-C3 |
| Fos in situ hybridization probe | Advanced Cell Diagnostics | 316921-C2 |
| Slc32a1 in situ hybridization probe | Advanced Cell Diagnostics | 319191-C3 |
| Software and algorithms |  |  |
| Image J | NIH | https://imagej.nih.gov/ij/ |
| GraphPad Prism | Dotmatics | Version 10.1.2 |
| MATLAB | MathWorks | https://github.com/djamesbarker/pMAT |

## Experimental methods

### Experimental animals

C57BL/6J mice were purchased from the Vital River Laboratory Animal Technology Co., Ltd. (Zhejiang, China). Calb1-IRES2-Cre-D (*Calb1^Cre^*) mice (strain #028532, B6;129S-*Calb1^tm2.1(cre)Hze^*/J) were kindly provided by Prof. Xiaohui Zhang (State Key Laboratory of Cognitive Neuroscience & Learning and IDG/McGovern Institute for Brain Research, Beijing Normal University). Mice were maintained in a specific pathogen-free (SPF) grade animal facility with controlled environments (23 ± 1°C, 50 ± 10% humidity, 12 h light/dark cycle). Mice were housed in groups of four to five per individually ventilated cages and given access to food and water *ad libitum*. Experimenters were blinded to the animals’ treatments and sample processing throughout the subsequent experimentation and analyses.

### Ethical approval and animal experimentation design

Animal experiment protocols were approved by the Animal Care Committee of the Southern University of Science and Technology (Shenzhen, China). The ARRIVE guidelines were followed when designing, performing, and reporting animal experimentation ^59^. Efforts were made to minimize the number of mice used. Mice used in the current study were randomly assigned to groups to maintain complete randomization. A minimum of five mice per group were used in the *in vivo* experiments. For tissue section staining, at least three mice were used per group. Mice at 3 months old, both male and female, were used in this study.

### Recombinant viruses

The following recombinant adenoviruses (rAAV) were used for stereotaxic injection into the brain

For fiber photometry recording of calcium *in vivo*: rAAV2/9-VGLUT2-DIO-GCaMP6s-WPRE (5.28 × 10¹² v.g./mL at 150 nL) was injected unilaterally into the DMH of *Calb1^Cre^* mice.

For chemogenetic modulation of DMH CALB1-positive and VGLUT2-positive (DMH^Calb1/Vglut2^) neurons: the rAAV2/9-VGLUT2-DIO-hM3D(Gq)-EGFP-WPRE (5.09 × 10¹² v.g./mL at 60 nL) (for activation of DMH^Calb1/Vglut2^ neurons) and rAAV2/9-VGLUT2-DIO-hM4D(Gi)-EGFP-WPRE (5.44 × 10¹² v.g./mL at 60 nL) (for inhibition of DMH^Calb1/Vglut2^ neurons) were bilaterally injected into the DMH of *Calb1^Cre^* mice, respectively. The control rAAV used was AAV2/9-VGLUT2-DIO-EGFP-WPREs (5.45 × 10¹² v.g./mL).

For chemogenetic modulation of DMH CALB1-negative and VGLUT2-positive (DMH^Calb1-/Vglut2+^) neurons: the rAAV2/9-VGLUT2-DO-hM3D(Gq)-mCherry-WPRE (5.01 × 10¹² v.g./mL at 60 nL) (for activation of DMH^Calb1-/Vglut2+^ neurons) and rAAV2/9-VGLUT2-DO-hM4D(Gi)-mCherry-WPRE (5.12 × 10¹² v.g./mL at 60 nL) (for inhibition of DMH^Calb1-/Vglut2+^ neurons) were bilaterally injected into the DMH of *Calb1^Cre^* mice, respectively. The control virus used was rAAV2/9-VGLUT2-DO-mCherry-WPREs (5.32 × 10¹² v.g./mL).

For optogenetic inhibition of DMH^Calb1/Vglut2^ neurons: rAAV2/9-VGLUT2-DIO-stGtACR2-EGFP-WPRE (5.22 × 10¹² v.g./mL at 200 nL) was bilaterally injected into the DMH of *Calb1^Cre^* mice. Bilateral injection of rAAV2/9-VGLUT2-DIO-EGFP-WPREs (5.45 × 10¹² v.g./mL, 200 nL) into the DMH of *Calb1^Cre^*mice served as the control.

For Diphtheria toxin A (DTA)-mediated neuronal deletion studies: rAAV2/9-VGLUT2-DIO-DTA-mCherry-WPRE-hGH polyA (5.05 × 10¹² v.g./mL at 50 nL) was bilaterally injected into the DMH of *Calb1^Cre^* mice. The control virus used was rAAV2/9-VGLUT2-DIO-EGFP-WPREs (5.45 × 10¹² v.g./mL).

For retrograde tracing of upstream neurons of DMH^Calb1/Vglut2^ neurons: the rAAV-EF1α-DIO-N2cG (2.5 × 10¹² v.g./mL) and rAAV-Vglut2-DIO-EGFP-T2A-TVA (5.0 × 10¹² v.g./mL) viruses were mixed in a 2:1 ratio and injected unilaterally into the DMH of *Calb1^Cre^* mice at a volume of 60 nL. Two weeks later, 60 nL of the RV-CVS-EnvA-ΔG-tdTomato (5.0 × 10^8^ IFU/mL) virus was also injected at the same site. The following week, mice were euthanized, and the brains were collected for sectioning and staining.

For ablating upstream neurons of DMH^Calb1/Vglut2^ neurons, a mixture of rAAV-EF1α-DIO-N2cG (2.5 × 10¹² v.g./mL) and rAAV-Vglut2-DIO-EGFP-T2A-TVA (5.0 × 10¹² v.g./mL) at a 2:1 ratio was bilaterally injected into the DMH of *Calb1^Cre^* mice at a volume of 80 nL. Meanwhile, AAV-nEF1α-FDIO-taCasp3-P2A-TEVp (5.1 × 10¹² v.g./mL) were bilaterally injected into the LSI, POA, and VLPAG, respectively. To ensure optimal viral coverage of the LSI and POA. Two injection sites (120 nL per site) per hemisphere were targeted along the dorsoventral axis. For the VLPAG, a single injection (60 nL per site) was delivered per hemisphere. Two weeks later, RV-CVS-EnvA-ΔG-mCherry-P2A-FlpO (2.26 × 10^8^ IFU/mL) was bilaterally injected into the DMH (80 nL per site).

### Brain stereotaxic injection

The method used was exactly as we previously described ^26^. Mice were injected subcutaneously with meloxicam (5 mg/kg) 30 minutes prior to surgery. Anesthesia was induced using 2.5% isoflurane with air (Product #R540, RWD Life Science, Shenzhen, China). The mice were then positioned on the stereotaxic apparatus under anesthesia with 1.5% isoflurane in air. Body temperature was maintained at 36.5 ± 0.5°C using a heating blanket (Product #TCAT-2DF, Harvard Apparatus, USA). After hair removal and sterilization of the scalp with povidone-iodine, a hole was drilled in the mouse’s skull with a dental drill (Product #78001, RWD Life Science), guided by a stereotaxic frame (Product #68861N, RWD Life Science) to target the bilateral or unilateral DMH (coordinates from Bregma: ML: ±0.1 mm, AP: −1.5 mm, and DV: −5.15 mm), POA (coordinates from Bregma: ML: ±0.25 mm, AP: 0.14 mm, and DV: −5.25 and -5.55 mm), LSI (coordinates from Bregma: ML: ±0.35 mm, AP: 0.62 mm, and DV: −3.22 and -3.70 mm), and VLPAG (coordinates from Bregma: ML: ±0.35 mm, AP: −4.36 mm, and DV: −2.7 mm). These coordinates were based on the mouse brain atlas.

Following the virus injection, the cranial hole was sealed with sterile bone wax, and the incision was closed using sterile surgical sutures. Meloxicam (5 mg/kg) was administered for the first three days after surgery, followed by a recovery period of at least three weeks for virus expression.

### Rectal thermometry

Rectal thermometry is a precise and straightforward method for measuring short-term T_core_ in rodents. The procedure was performed as previously described ^26^. Mice were anesthetized with 1.5% isoflurane in air, and a thermocouple probe (KPS-TT-T-36-SLE, KAIPISEN, China), covered with Vaseline, was gently inserted into the rectum to a depth of 2 cm. The probe wire was secured to the tail using medical waterproof fabric, and the mice were allowed to recover. The flexible, plastic-covered thermocouple wires provided good deformability, allowing the mice to move freely in their home cage while connected to the thermometer (DC700 multichannel temperature recorder, DCUU, Pumei, China) via the long wire. After a 30-minute adaptation period, mice with the rectal thermoprobe exhibited no signs of discomfort or resistance during rectal temperature monitoring.

### Hypothermia evoked by surface cooling

Mice in their home cage, equipped with implanted rectal temperature measurement probes, were sprayed with 1 mL of 70% ethanol on their dorsal skin to wet the fur. Next, ice-cold water was evenly dripped onto the dorsal skin to lower the T_core_, and the mouse’s T_core_ was monitored. Within approximately 10 minutes, the T_core_ was reduced to 33°C, and the total volume of ice-cold water used was recorded. Food was withheld during the cooling process. Throughout the experiment, each mouse experienced only one T_core_ reduction episode.

### Intramuscular EMG recording and signal analysis

Intramuscular EMG recordings were performed using a Pinnacle EMG head mount (Product #8201-SS, Pinnacle, USA) following the manufacturer’s guidelines. Prior to surgery, the mouse received a subcutaneous injection of meloxicam (5 mg/kg) and was anesthetized with 1.5% isoflurane in air. The mouse was then positioned on a stereotaxic apparatus, and body temperature was maintained at 36.5 ± 0.5°C using a heating blanket. After hair removal and sterilization with povidone-iodine, a section of the mouse’s scalp was removed, and the EMG headmount was securely attached to the skull using four screws (Product #8209, #8212, Pinnacle, USA). Two flexible electrode wires were inserted deep into the dorsal cervical skeletal muscles, and the headmount was further stabilized with dental cement. For the first 3 days post-surgery, meloxicam (5 mg/kg) was administered, followed by a 4-day recovery period. EMG signals, amplified (×100) and filtered (10-1000 Hz), were recorded from freely moving mice using the data conditioning and acquisition system (Product #8200-K3-SL/SE, Pinnacle, USA). Simultaneous EMG and video recordings of mouse movement activity were performed at normal body temperature and under hypothermic conditions. Video recordings enabled identification of movement and rest states, allowing the selection of resting-state EMG signals for analysis.

EMG signal analysis was performed using established methods as previously described ^15, 17, 60^. The raw EMG signal was sequentially processed to eliminate DC offset, power-line noise (50 Hz), environmental noise from variable-frequency drives and motors, and slow drift and movement artifacts ^61, 62^. Then, the EMG signal was quantified by calculating the mean root-mean-square (RMS) of total power in the 20–150 Hz frequency band per second, using a custom MATLAB script to assess cervical muscle mechanical activity. At an ambient temperature of 25°C, the mean RMS of the EMG signal from the neck muscles of a resting mouse indicated muscle tone ^15, 17^. After reducing the mouse’s T_core_ to below 35°C with an ice-cold water spray, the burst of rhythmic muscle contractions recorded at rest was identified as overt shivering.

### Electrical lesion of the DMH

Following the same procedure as virus injection, a conductive-tip electrode (KedouBC, China) was stereotaxically implanted into the DMH. An electrical lesion was induced by passing a 0.4 mA current for 20 seconds using a Lesion Making Device (Product #53500, Ugo Basile, Italy). Meloxicam (5 mg/kg) was administered for the first 3 days post-surgery, followed by a 4-day recovery period. For the sham-operated group of mice, all procedures were the same except for the electrical lesion. To confirm the lesioned area, mice were transcardially perfused with 0.01M phosphate-buffered saline (PBS), followed by 4% paraformaldehyde under deep anesthesia. After brain fixation, dehydration, and sectioning, the brain sections (16 μm in thickness) containing the lesion site were stained with crystal violet (0.5% in 20% alcohol solution with 660 μL acetic acid) for 8 minutes. The sections were then washed twice with 100% alcohol and immersed in xylene. Finally, the slides were mounted with neutral resin, and brain images were captured using a microscope (Tissue FAXS Plus, Meyer Instruments, Inc., USA).

### Retrograde trans-synaptic pseudorabies virus (PRV) tracing

Mice were anesthetized with 1.5% isoflurane in air and placed on a heated pad to maintain body temperature. After fur removal and skin sterilization with povidone-iodine, the dorsal cervical skeletal muscles were exposed. Three injection sites were created at a depth of 1.5 – 2.0 mm into the muscles using a microsyringe (Hamilton 600, USA). At each site, 0.5 μL of either PRV-CAG-EGFP (2.0 × 10LJ PFU/mL) was injected at a rate of 300 nL/min using a pump. Following each injection, the syringe was left in place for 3 minutes to ensure complete virus delivery. After the PRV injection, the wound was sutured, and the mice were allowed a 4-day recovery period. Subsequently, the mice were euthanized via an intraperitoneal injection of sodium pentobarbital (70 mg/kg). Fresh brain tissue was collected for RNA and protein analysis, and transcardial perfusion fixation was performed for histological sectioning.

### DMH neuron RNA-seq

Retrograde trans-synaptic PRV tracing was performed by injecting 1.5 μL of PRV-CAG-EGFP into the dorsal cervical skeletal muscle. Four days post-injection, mice were anesthetized with sodium pentobarbital (70 mg/kg, i.p.) and sacrificed for brain extraction. The brains were immediately transferred to ice-cold Hibernate™-A medium (A1247501, Gibco). Using a brain matrix, 1 mm-thick coronal sections were prepared, and the DMH nucleus was identified under a stereomicroscope and carefully dissected.

The excised DMH tissue was minced in Hibernate™-A medium containing papain (2 mg/mL) and DNase (10 μg/mL), then incubated at 37°C with gentle shaking at 180 rpm for 30 minutes. After enzymatic digestion, the tissue suspension was triturated using a pipette and filtered through a 70 μm cell strainer. The filtrate was then centrifuged at 300 × g for 5 minutes at 4°C to collect the cells, which were subsequently resuspended in Hibernate™-A medium for downstream RNA sequencing analysis.

EGFP-positive and EGFP-negative cells were separated using a flow cytometer (BD FACSAria SORP, BD Biosciences). Collected cells were centrifuged at 300 × g for 5 minutes at 4°C. EGFP-positive neurons were immediately lysed and frozen for RNA sequencing. EGFP-negative cells underwent additional purification using the Adult Brain Neuron Isolation Kit (130-107-677, Miltenyi Biotec China) combined with magnetic bead sorting (130-042-201, 130-042-102, Miltenyi) to isolate neuronal populations, and were subsequently processed for RNA sequencing.

The experiment comprised two groups: EGFP-positive and EGFP-negative neurons, each with three biological replicates. Samples were submitted to BGI Genomics (Shenzhen, China) for RNA-seq. Data analysis was performed using BGI’s Dr. Tom platform to identify differentially expressed genes between retrogradely trans-synaptically labeled DMH EGFP-positive neurons and non-labeled neurons. This approach facilitated the characterization of neuronal subtypes and the identification of potential molecular markers for DMH neurons involved in shivering regulation.

### Fiber photometry setup and recording

Two weeks following viral microinjection, optical fibers and EMG headmounts were surgically implanted into the mice. Fiber implantation was conducted using a rotational digital stereotaxic frame for mice (Model 69100, RWD Life Science, Shenzhen, China). The sagittal axis was rotated by ±27.5° to define the implantation coordinates relative to bregma: mediolateral (ML), ±0.26 mm; anteroposterior (AP), −1.5 mm; dorsoventral (DV), −5.16 mm. A single optical fiber (outer diameter, 200 μm; numerical aperture, 0.37; Thinker Tech Nanjing Biotech, China) was unilaterally implanted at 0.01 mm above the viral injection site.

Following fiber implantation and initial fixation with tissue adhesive, the EMG headmount was installed and secured with dental cement. After a one-week recovery, the headmount was connected to the electrophysiology recording system. Neuronal calcium signals were recorded using a commercial two-channel fiber photometry system (Thinker Tech Nanjing Biotech, China), which included a 405 nm LED for isosbestic control and a 470 nm LED for fluorescence excitation. Before recording EMG and calcium signals simultaneously, mice were allowed to acclimate to their home cages for 30 minutes.

To characterize calcium signal dynamics during T_core_ reduction to 33-36°C, the experimental protocol was conducted as follows. First, calcium signals were continuously acquired for 5 min under basal conditions to record baseline activity, after which data acquisition was temporarily halted. Second, ice-cold water was used to sequentially regulate the T_core_ to 35 °C, 34 °C, and 33 °C. At each target temperature (with mice kept stationary), calcium signals were recorded for 5 min.

### Fiber photometry data analysis

Initial fiber photometry data preprocessing was performed using custom MATLAB scripts, modified from the open-source pMAT (Photometry Modular Analysis Tool) package developed by the Barker Lab (https://github.com/djamesbarker/pMAT), following previously established protocols for raw trial data processing ^63^. Subsequent fluorescence signal processing and quantitative analysis were performed according to well-established standard pipelines for fiber photometry recordings ^64^. Briefly, raw data from the functional signal channel (470 nm) and isosbestic control channel (405 nm) were extracted separately. Both channel signals were then smoothed using the locally weighted scatterplot smoothing (Lowess) algorithm, which performs local linear regression. Signal amplitude normalization was further achieved via the MATLAB polyfit function. Specifically, raw fluorescence traces were corrected against isosbestic control signals to eliminate photobleaching artifacts and minimize non-specific signal fluctuations:

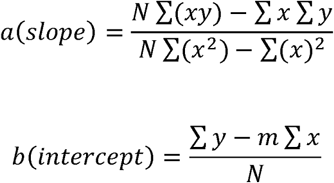

N represents the number of subjects, x denotes the control channel, and y refers to the signal channel. The resulting slope and intercept are then used to generate a scaled control channel. This process is visualized in a MATLAB figure titled ’Signal vs. Fitted Control:

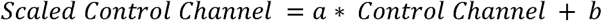

Subsequently, the delta F/F (ΔF/F) is generated by subtracting the fitted control channel from the signal channel:

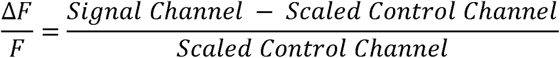

Finally, the signals were transformed into robust z-scores using the following formula:

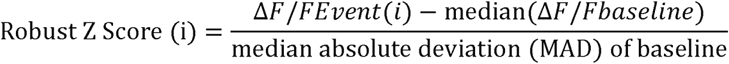

### Chemogenetic manipulation of DMH neurons

Chemogenetic manipulation of DMH neurons was performed as previously described experimental protocols ^26, 63^. All mice were habituated to the experimental environment for at least 30 min before behavioral and physiological recordings. Baseline T_core_ and EMG signals were synchronously acquired throughout a 30-min resting session. Following baseline recording, *Calb1^Cre^*mice with DMH expression of DIO-hM4D(Gi), DIO-hM3D(Gq), or control virus received an intraperitoneal injection of either sterile saline or CNO (3 mg/kg, dissolved in saline). One hour after injection, ice-cold water was sprayed onto the mouse body surface to gradually reduce T_core_ to 33 °C, at which point water spraying was terminated. Continuous data acquisition was maintained until T_core_ spontaneously recovered to baseline levels.

### Optogenetic manipulation of DMH neurons

Two weeks following viral microinjection, mice underwent surgical implantation of optical fibers and EMG headmounts. Fiber implantation was conducted using a rotational digital stereotaxic frame for mice (Model 69100, RWD Life Science). The sagittal axis was rotated by ±27.5° to locate the target implantation site, with stereotaxic coordinates defined relative to bregma as follows: ML, ±0.26 mm; AP, −1.5 mm; DV, −5.16 mm. Bilateral optical fibers (200 μm outer diameter, Model R-FOC-L200C-39NA, RWD Life Science) were implanted 0.01 mm above the viral injection sites. After positioning the optical fiber and preliminary fixation with tissue adhesive, the EMG headmount was affixed and secured with dental cement.

One week after implantation, the EMG headmount was coupled to the recording apparatus. Optical fibers were connected to a Smart Light Source (B1402, Inper, China) for optogenetic stimulation. All mice were habituated to their home cages for a minimum of 30 min prior to behavioral and physiological recordings. Bilateral yellow light stimulation (589 nm, constant power of 15 mW) was delivered continuously for 2 min, followed by a 3-min laser-off interval. EMG signals were recorded synchronously at baseline room temperature and during ice-cold water-induced core body temperature reduction to 33 °C.

### Heart rate and pulse distention assessment

One day after cervical hair removal with depilatory cream, a sensor clip was attached to the cervical region of mice housed in their home cages. Following a 30-min habituation period, heart rate and pulse distention were measured using a MouseOx® Plus Pulse Oximeter (STARR Life Science, USA). Baseline physiological signals were continuously recorded for 10 min, after which mice received an intraperitoneal injection of CNO (3 mg/kg). Thirty minutes post-injection, heart rate and pulse distention were recorded for an additional 10 min during chemogenetic activation or inhibition of DMH^Calb1/Vglut2^ neurons.

### Infrared imaging of body temperature

Prior to infrared imaging conducted at a constant room temperature of 25 ± 0.5 °C, all mice were habituated in their home cages for at least 30 min. Infrared thermal images were acquired at two time points: before experimental stimulation and 30 min following CNO administration. Image collection was performed using a handheld infrared camera, and all acquired thermal images were subsequently analyzed using supporting software.

### Determination of brown fat tissue temperature (T_BAT_)

Mice were anesthetized with 1.5% isoflurane, and the interscapular brown adipose tissue (iBAT) was surgically exposed under sterile conditions as previously described ^26, 65, 66^. A sterile thermocouple probe was inserted into the iBAT parenchyma to measure temperature. The thermocouple wire was firmly fixed in place, and the surrounding skin was sealed with 3M™ Scotch-Weld™ Surface Insensitive Instant Adhesive SI Gel (3M, USA) prior to surgical suture. Real-time iBAT temperature was continuously recorded at room temperature using a DC700 multichannel temperature recorder (PUMEI, China) connected to the implanted thermocouple probe. Data were collected and analyzed using GraphPad (version 9.0.0, USA).

### Electrophysiological recordings in acute brain slices

Acute brain slice preparation was performed following previously established protocols ^49, 67, 68^. Mice were deeply anesthetized via intraperitoneal injection of sodium pentobarbital (70 mg/kg) and subsequently euthanized for rapid brain dissection. Freshly isolated whole brains were immediately transferred to ice-cold oxygenated cutting buffer (pH = 7.4 ± 0.1) containing the following components (in mM): 124 NaCl, 2.5 KCl, 1.25 NaHPO, 25 NaHCO, 2 CaCl, 2 MgSO, 15 D-glucose, and 220 sucrose, continuously aerated with 95% O_2_ and 5% CO_2_. Coronal brain slices (350 μm thickness) containing the DMH were sectioned using a vibratome (VT1120S, Leica Systems). Obtained slices were rinsed twice with standard artificial cerebrospinal fluid (aCSF; pH = 7.4 ± 0.1), consisting of (in mM): 124 NaCl, 2.5 KCl, 1.25 NaH_2_PO_4_, 25 NaHCO_3_, 2 CaCl_2_ 2 MgSO_4_, and 15 D-glucose. Slices were then transferred to aCSF continuously bubbled with 95% O_2_/5% CO_2_, incubated in a 36 °C water bath for 30 min, and subsequently maintained at room temperature for at least 1 h prior to electrophysiological recording. For recordings, slices were placed in a recording chamber (RC26G, Warner Instruments, USA) mounted on the motorized x–y stage of an upright microscope (BX51W, Olympus, USA). During data acquisition, the slice chamber was continuously perfused with oxygenated aCSF at a constant flow rate of 10 mL/min. The acquisition frequency was 20.0 kHz, and the filter was set to 2.9 kHz. The number of action potentials (AP) was quantified from neuronal spiking using Mini-analysis (Synaptosoft Inc., USA).

To evaluate the thermosensitive properties of DMH^Calb1/Vglut2^ neurons, we performed bilateral microinjection of the AAV2/9-VGLUT2-DIO-EGFP-WPRE vector into the DMH region of *Calb1^Cre^* mice to label DMH Calb1/Vglut2-expressing neurons. Patch-clamp recordings were conducted on EGFP-labeled DMH^Calb1/Vglut2^ neurons in aCSF-perfused acute brain slices. The bath temperature within the recording chamber was precisely regulated at 33 °C, 35 °C, 36 °C, or 39 °C using an inline solution heater (SH-27B, Warner Instruments, USA) coupled with a temperature controller (TC-324C, Warner Instruments, USA). Recording pipettes were filled with an internal solution containing (in mM): 128 potassium gluconate, 10 NaCl, 10 HEPES, 0.5 EGTA, 2 MgCl_2_, 4 Na_2_ATP, and 0.4 NaGTP. To block the excitatory and inhibitory synaptic transmissions to DMH^Calb1/Vglut2^ neurons, a cocktail of blockers containing (in µM): 20 DNQX, 50 D-AP5, and 20 gabazine ^36^. A salt bridge containing 3 M KCl was used to ground the chamber and correct liquid junction potentials, as previously described ^37, 69^.

Spontaneous AP firings were recorded for 2 min at each target temperature, with only a single neuron recorded per brain slice. For temperature ramp assays, AP firing activity was first acquired at 36 °C, followed by sequential recordings at 33 °C and 39 °C, with a 5-min stabilization interval between each temperature shift.

As reported in prior studies, neurons were defined as cold-sensitive when exhibiting a thermal coefficient (TC) of ≤ −0.6 impulses·s⁻¹·°C⁻¹ ^26, 70^. The TC quantifies the responsiveness of neuronal firing rate to temperature fluctuations. In cold-sensitive neurons, a negative TC indicates elevated firing activity at lower temperatures, consistent with established classification criteria.

To determine the involvement of K^+^ channels in low-temperature-induced activation of DMH^Calb1/Vglut2^ neurons, we first recorded spontaneous AP at 36°C and 33°C in aCSF supplemented with DNQX (20 μM), D-AP5 (50 μM), and gabazine (20 μM). K⁺ currents were then measured in aCSF containing 1 μM TTX and 50 μM CdCl₂ to eliminate voltage-gated Na⁺ and Ca²⁺ currents. At a holding potential of −70 mV, outward K⁺ currents (*I*_K_ and *I*_A_) were evoked by 500-ms voltage steps (−90 mV to 20 mV, 10 mV increments) after a 300-ms pre-pulse to −110 mV. A 100-ms pre-step to −10 mV was then introduced to inactivate *I*_A_ and isolate *I*_K_. *I*_A_ was derived via offline subtraction of currents recorded with and without the −10 mV pre-step in the same cell as previously described ^49^. The calculation method for *I*_A_ inactivation time constant was adopted from established published procedures ^46, 71^.

### Cannula implantation and synaptic blocker infusion

Stereotaxic cannula implantation for localized brain drug delivery was performed according to our previously established protocols ^26^. Briefly, mice received a subcutaneous injection of meloxicam (5 mg/kg) 30 min prior to surgery for preoperative analgesia. General anesthesia was initially induced with 2.5% isoflurane in ambient air, after which mice were secured on a stereotaxic frame and maintained under anesthesia with 1.5% isoflurane in air throughout the surgical procedure. A heating blanket was used to sustain the core body temperature of mice at 36.5 ± 0.5°C during surgery. Following scalp hair removal and disinfection with povidone-iodine, bilateral cranial holes were drilled using a dental drill. To target the precise implantation site, the sagittal axis of the stereotaxic apparatus was rotated ±27.5°, with stereotaxic coordinates defined relative to the bregma: ML = ±0.26 mm, AP = −1.5 mm, and DV = −5.16 mm. A bilateral guide cannula (62010, RWD Life Science) was subsequently implanted into the DMH. After temporary fixation of the guide cannula with tissue adhesive, an EMG headmount was installed and firmly secured with dental cement. All experimental mice underwent daily health monitoring for seven consecutive days postoperatively.

On the day of testing, mice were gently restrained, and the EMG headmount was connected to electrophysiological recording equipment. An injector needle (62204, RWD Life Science) was inserted into the pre-implanted guide cannula, and the needle was connected to a syringe infusion pump (R462, RWD Life Science) via plastic tubing. Mice were allowed to freely explore the experimental arena for 20 min, during which baseline EMG and T_core_ recordings were acquired.

Following baseline acquisition, bilateral intra-DMH infusion of the synaptic blocker cocktail was performed at a constant rate of 50 nL/min for a total volume of 50 nL per hemisphere. After 25 min of uninterrupted EMG and T_core_ monitoring post the first infusion, a second identical infusion was delivered. Five minutes after the completion of the second infusion, ice water stimulation was applied to elicit T_core_ reduction, while EMG signals were recorded continuously throughout the entire stimulation and post-stimulation phase.

To block upstream synaptic input to DMH^Calb1/Vglut2^ neurons in mice, we followed a protocol previously used for rats ^72^. However, pilot studies in mice showed that 5 mM of DNQX and D-AP5, used in rats as described, were toxic to mice. Subsequent experiments employed a cocktail of synaptic transmission blockers containing DNQX, D-AP5, and gabazine, all at 0.1 mM in saline, which showed no toxic effects in mice. The cocktail was injected bilaterally into the DMH of C57BL/6J mice, delivering 50 nL per side at 50 nL/min. After 25 min of continuous post-infusion EMG and T_core_ recordings, a second identical infusion was carried out. Five min after completing the second infusion, an ice-water spray was applied to induce a reduction in T_core_, and EMG signals were continuously recorded during both the stimulation and post-stimulation periods.

### Immunofluorescence staining

Histological procedures were conducted according to previously described methods ^26, 63^. Briefly, mice were deeply anesthetized via intraperitoneal injection of sodium pentobarbital (70 mg/kg). Under deep anesthesia, transcardial perfusion was first performed with 0.01 M PBS, followed by freshly prepared 4% paraformaldehyde (PFA). Harvested brain tissues were post-fixed overnight in 4% PFA at 4 °C. For cryoprotection, brain samples were sequentially dehydrated in 10% and 30% sucrose solutions. Tissue sections 20 μm thick were then cut using a cryostat microtome (CM1950, Leica, Germany) for subsequent immunofluorescence staining. After cardiac perfusion with 0.01 M PBS in mice, the brains were quickly removed, embedded in Optimal Cutting Temperature compound, and rapidly frozen on dry ice. The brains were then sectioned at 10 μm using a cryostat for immunofluorescence or FISH analysis.

Immunofluorescence staining was performed according to established laboratory protocols ^26, 49, 67^. Briefly, brain sections were subjected to antigen retrieval as previously reported ^26^, after which sections were blocked with 5% BSA for 1 h to eliminate non-specific antibody binding. For primary antibody incubation, sections were incubated overnight at 4 °C in a humidified chamber with primary antibodies diluted in 3% BSA. The primary antibodies used included mouse anti-c-Fos (ab208942, Abcam, USA, 1:500), rabbit anti-c-Fos (226008, Synaptic Systems, Germany, 1:500), and rabbit anti-Calbindin-D28k (14479-1-AP, Proteintech, China, 1:500). Following thorough PBS washing, sections were incubated with corresponding secondary antibodies, including Alexa Fluor 594-conjugated goat anti-mouse IgG (ab150116, Abcam, USA, 1:500), Alexa Fluor 594-conjugated goat anti-rabbit IgG (ab150080, Abcam, USA, 1:500), and Alexa Fluor 488-conjugated goat anti-rabbit IgG (ab150077, Abcam, USA, 1:500). After secondary antibody incubation, slides were mounted with DAPI-containing anti-fade mounting medium (Fluor Shield, H-1200-10, Vector Laboratories, USA) and covered with coverslips for subsequent imaging analysis.

### RNAscope fluorescence in situ hybridization (FISH)

FISH was conducted using the RNAscope Multiplex Fluorescent Assays V2 (323110, Advanced Cell Diagnostics, USA) in strict accordance with the manufacturer’s standard protocols. Target-specific probes for *Slc17a6* (1170921-C1), *Egfp* (538851-C2), *td-Tomato* (317041-C2), *Calb1* (428431-C3), *Fos* (316921-C2), and *Slc32a1* (319191-C3) were all obtained from Advanced Cell Diagnostics Inc. Fluorescent images were acquired using multiple microscopic systems, including a Tissue FAXS Plus fluorescence microscope (Meyer Instruments, Inc., USA), a LSM980 confocal microscope (Zeiss, Germany), and an Axio M2 fluorescence microscope (Zeiss, Germany). All acquired images were quantitatively analyzed using ImageJ software as we previously described ^26^.

### Quantification and statistical analysis

All statistical analyses were performed using GraphPad Prism software (Version 10.1.2, San Diego, CA, USA). All data are presented as the mean ± standard deviation (SD) or the mean ± standard error of the mean (SEM), as indicated in the text. The Shapiro-Wilk test was applied to assess data distribution prior to statistical testing, guiding the selection of parametric or nonparametric analytical approaches. For two-group comparisons, unpaired or paired t-tests were used for normally distributed data, whereas the Mann-Whitney U test was used for non-normally distributed data. Multiple-group comparisons were conducted via one-way or two-way analysis of variance (ANOVA), followed by Tukey’s or Sidak’s *post hoc* tests for pairwise multiple comparisons. Neuronal firing rate data were analyzed using two-way repeated-measures (RM) ANOVA with Tukey’s or Sidak’s *post hoc* testing. Nonlinear regression or simple linear regression analysis was utilized to assess the correlations among T_core_, EMG power, and calcium signal activity. Detailed statistical information, including sample size (n), data precision metrics, statistical test types, and significance criteria, was provided in individual figure legends. A value of p < 0.05 was defined as the threshold for statistical significance.

### Reporting summary

Further information on research design is available in the Nature Portfolio Reporting Summary linked to this article.

## Data availability

All rAAV constructs generated from this study are available from the lead contact upon request. Further information and requests for resources and reagents should be directed to and will be fulfilled by the Lead Contact, Sheng-Tao Hou.

## Code availability

All data reported in this paper will be shared by the lead contact upon request. The MATLAB code used for EMG analysis has been deposited on GitHub and is available under the following link: https://github.com/ASAI243/EMG.git. Any additional information required to reanalyze the data reported in this paper is available from the lead contact upon request.

## Acknowledgments

We would like to thank the SUSTech Animal Facility for animal care and the SUSTech Core Research Facilities for their support in imaging acquisition. We are also deeply grateful to Professor Xiaohui Zhang (State Key Laboratory of Cognitive Neuroscience & Learning, IDG/McGovern Institute for Brain Research, Beijing Normal University) for providing the *Calb1^Cre^* mice and to Dr. Ju Jun for providing initial training in patch-clamping techniques to one of the authors. Financial support for STH was provided by the National Natural Science Foundation of China (32371029), the Shenzhen Medical Research Fund (B2301001), the Natural Science Foundation of Guangdong Province (2023B0303040004), and the Shenzhen-Hong Kong Institute of Brain Science-Shenzhen Fundamental Research Institutions (2023SHIBS0002). STH is a member of the Guangdong Innovation Platform for Translational Research in Cerebrovascular Diseases and the SUSTech-UQ Joint Centre for Neuroscience and Neural Engineering (CNNE). Additionally, STH holds the position of Pengcheng Peacock Plan (A) Distinguished Professor.

## Author Contributions

SZ and HZ performed electrode implantation, neuron isolation, virus injections, immunofluorescence staining, FISH staining, histopathological evaluation, recordings for body temperature, BAT temperature, EMG signal, data analysis, and wrote the first draft of the manuscript; ZTH and YXW performed brain slice electrophysiology and data analysis; SYW performed virus injection, calcium signal recording, and data analysis; WH performed virus injection; DHL and LXH performed mice feeding and electronic lesion of DMH; RBL performed flow cytometry sorting; SFC performed mice maintenance. ZYZ and XYY performed EMG and fiber photometry data analysis; STH conceived the idea, secured funding, designed the experiments, analyzed the results, revised the first draft, and wrote the final version of the manuscript.

## Additional information

**Extended data** is available for this paper at https://doi.XXXXX

**Supplementary information** The online version contains supplementary material available at https://doi.XXX.

**Correspondence and requests for materials** should be addressed to Prof. Sheng Tao Hou.

