## Extended Data for "Shivering thermogenesis driven by hypothermia-sensitive neurons in the dorsomedial hypothalamus"


**Running title**: DMH neurons drive shivering

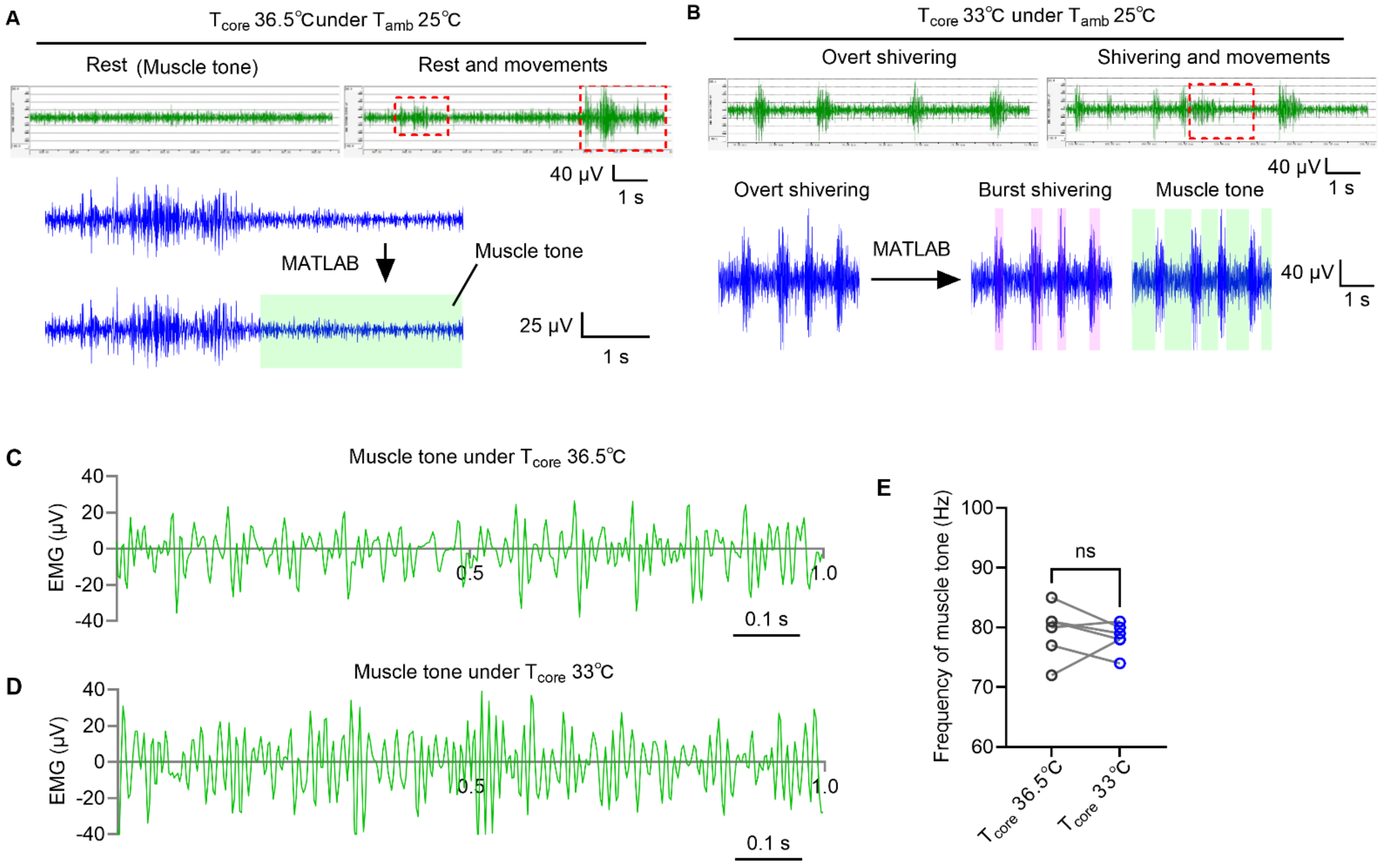


**Extended Data Fig. 1: Cervical muscle EMG analysis in freely moving mice at $T_{core}$ of 36.5 °C and 33 °C.**

(A) Representative cervical EMG traces recorded at an ambient temperature ($T_{amb}$) of 25 °C in mice with $T_{core}$ = 36.5 °C. Baseline resting EMG fluctuated within ±30 μV, while movement elevated signal amplitudes beyond this threshold (red dashed boxes). Custom MATLAB scripts isolated resting muscle tone for quantification via mean root-mean-square (RMS) power analysis.

(B) Representative cervical EMG traces after ice-cold water-induced reduction of $T_{core}$ to 33 °C at $T_{amb}$ = 25 °C. Hypothermia produced rhythmic shivering bursts; locomotion disrupted these rhythmic signals, generating movement-associated EMG activity (red dashed boxes). MATLAB segmentation distinguished shivering bursts (>30 μV) from basal muscle tone (±30 μV), with both signals assessed using mean RMS.

(C, D) Representative isolated muscle tone traces at $T_{core}$ = 36.5 °C (C) and $T_{core}$ = 33 °C (D).

(E) Quantitative comparison of muscle tone frequency derived from panels C and D.

Data are presented as mean ± SD, with n = 6 mice per group. Statistical comparisons were performed using paired t-tests for panel E, with n.s. indicating not significant ($p > 0.05$).

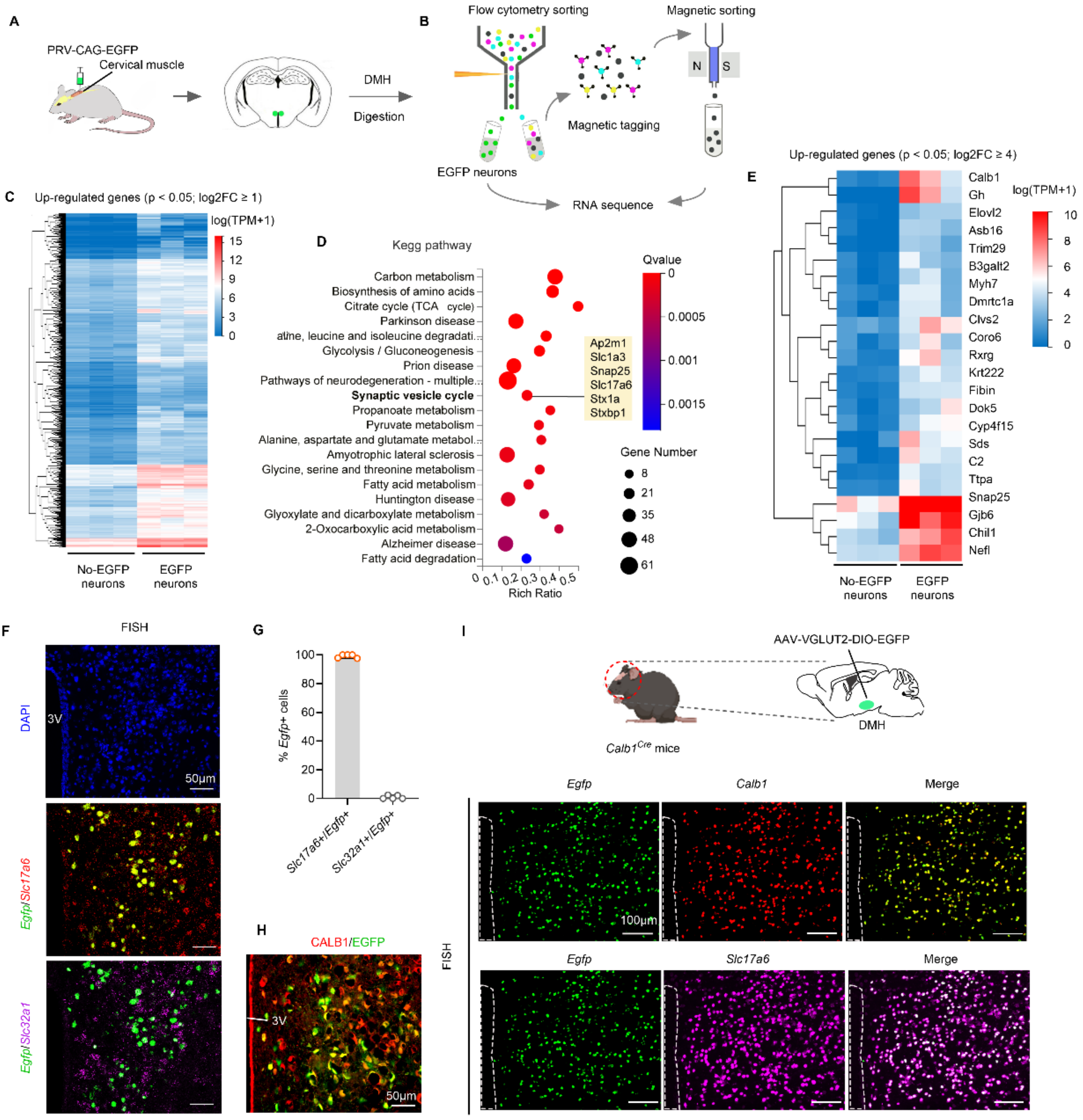


**Extended Data Fig. 2: RNA-seq profiling and bioinformatic analysis to identify DMH neurons retrogradely traced from cervical muscles.**

(A, B) Schematic workflow for retrograde PRV-CAG-EGFP tracing from the posterior cervical muscles to label DMH neurons. EGFP$^{+}$ and EGFP$^{-}$ neuronal populations were isolated via flow cytometry sorting for subsequent RNA sequencing.

(C, D) Heatmap and KEGG pathway enrichment analysis of upregulated genes identified in panel B; n = 3 biological replicates per group.

(E) Heatmap displaying the top 22 significantly upregulated genes ($q < 0.05$, $\log_2$ fold change ≥ 4) from the dataset in panel C.

(F) Representative FISH micrographs of DMH neurons retrogradely labeled with PRV-CAG-EGFP. $EGFP^+$ DMH neurons colocalized with *Slc17a6* (VGLUT2) but not *Slc32a1* (VGAT). Scale bar = 50 μm.

(G) Quantification of the percentage of $EGFP^+$ DMH neurons co-expressing *Slc17a6* or *Slc32a1*, corresponding to panel F.

(H) Representative immunofluorescence images confirming CALB1 expression in PRV-labeled $EGFP^+$ DMH neurons traced from cervical muscles. Scale bar = 50 μm.

(I) Experimental schematic for targeted labeling of $DMH^{Calb1/Vglut2}$ neurons via AAV-Vglut2-DIO-EGFP delivery into the DMH of $Calb1^{Cre}$ mice. Representative FISH images verify co-expression of EGFP with *Calb1* and *Slc17a6* in targeted DMH neurons. Scale bar = 100 μm.

Data were presented as means ± SD, with n = 5 mice.

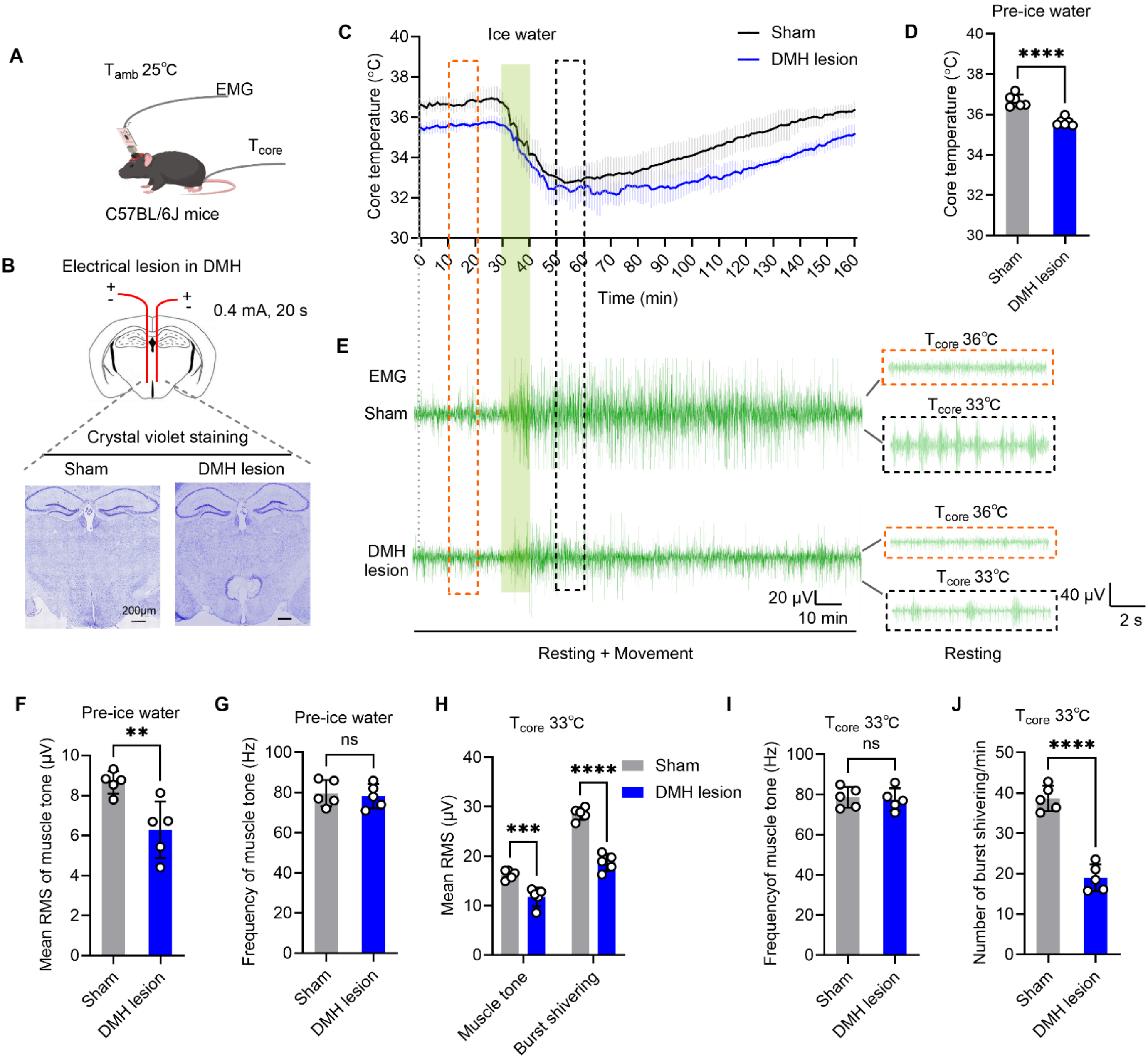


**Extended Data Fig. 3: DMH ablation attenuates hypothermia-evoked shivering.**

(A) Schematic for concurrent $T_{core}$ and EMG monitoring in freely moving mice following DMH lesion.

(B) Schematic and Cresyl violet verifying the site of electrical DMH lesions. Scale bar, 200 μm.

(C) Continuous recordings of $T_{core}$ of DMH-lesioned (blue trace) and sham-operated (black trace) mice during ice-water cooling to 33 °C; green bar denotes cooling duration.

(D) Quantitative analysis of $T_{core}$ changes defined in the time windows shown in panel C (orange dashed line).

(E) Representative cervical muscle EMG recordings from awake mice.

(F, G) Quantification of resting muscle tone EMG power (F) and frequency (G) before cold stimulation, corresponding to the orange dashed intervals in (E).

(H–J) Quantification of EMG power for muscle tone and burst shivering (H), muscle tone EMG frequency (I), and the number of burst shivering (J) when mouse $T_{core}$ reached 33°C, 10 minutes after ice-cold water spray, as shown in panel E (black dashed line).

Data were expressed as mean ± SD, with n=5 mice. Unpaired t-tests were applied for panels D, F, G, I, J; one-way ANOVA with Tukey's *post-hoc* test was used for panel H. **$p < 0.01$, ***$p < 0.001$, ****$p < 0.0001$. Scale bar = 200 μm.

Cre-on mice

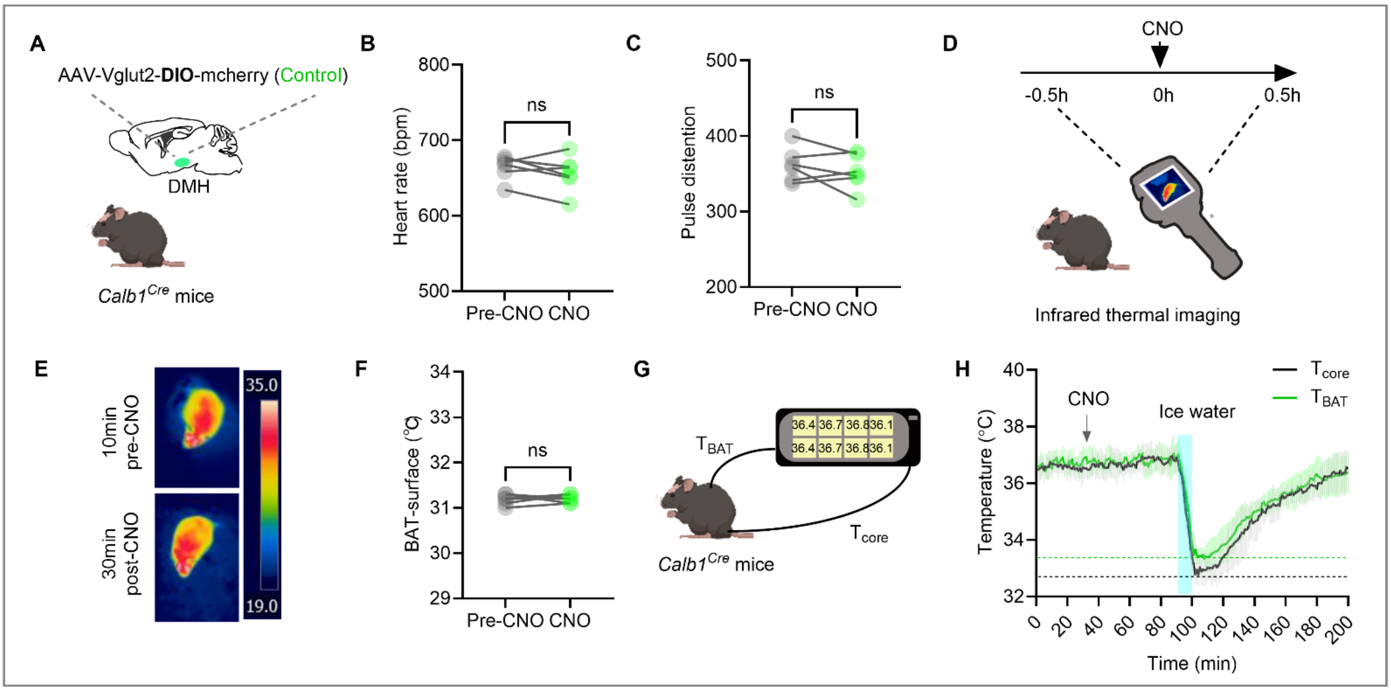


Cre-off mice

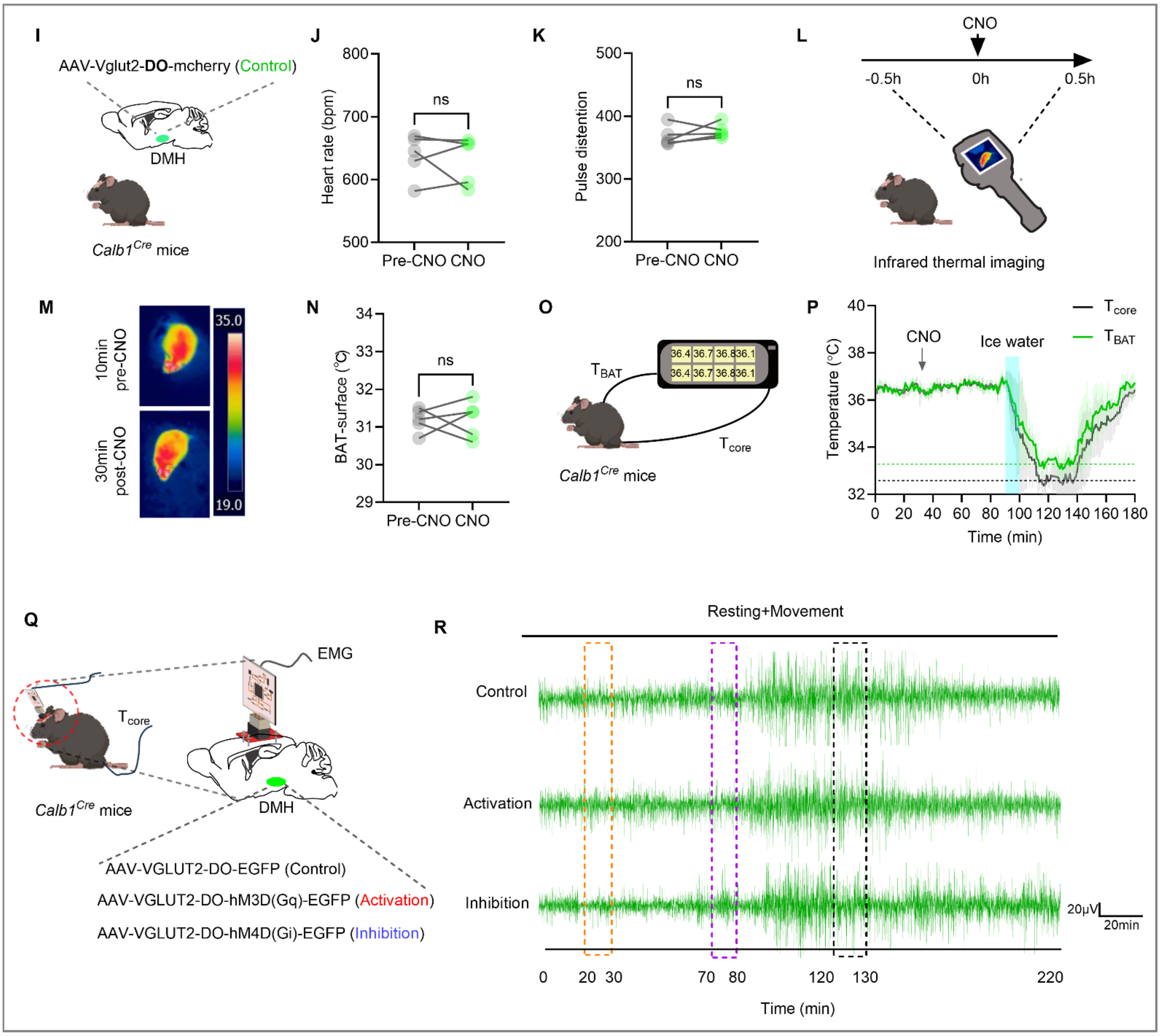

**Extended Data Fig. 4: Cre-on and Cre-off manipulation of Calb1$^-$/Vglut2$^+$ neurons in DMH and their effect on non-shivering and shivering thermogenesis.**

(A) Schematic of Cre-dependent DIO control virus delivery to the DMH of *Calb1$^{Cre}$* mice to target DMH Calb1$^+$/Vglut2$^+$ neurons (Cre-on configuration).

(B, C) Quantification of heart rate and pulse distention assessed 10 min before and 30 min after CNO injection in freely behaving animals.

(D–F) Infrared thermography workflow (D), representative dorsal thermal images (E), and quantitative interscapular $T_{BAT}$ analysis (F) at 10 min pre-CNO and 30 min post-CNO treatment.

(G) Schematic for simultaneous $T_{core}$ and $T_{BAT}$ recordings using implanted thermocouples in awake mice.

(H) Time-course traces of $T_{core}$ (black) and $T_{BAT}$ (green) with or without ice-cold water cooling to reduce $T_{core}$ to 33 °C following CNO administration.

(I) Schematic for Cre-off DO control virus injection into the DMH of *Calb1$^{Cre}$* mice to label DMH Calb1$^-$/Vglut2$^+$ neurons.

(J, K) Post-CNO quantification of heart rate and pulse distention, compared between 10 min pre-treatment and 30 min post-treatment.

(L–N) Infrared imaging schematic (L), representative dorsal thermal photographs (M), and statistical $T_{BAT}$ comparison (N) before and after CNO delivery.

(O) Schematic illustrating concurrent $T_{core}$ and $T_{BAT}$ recordings via thermocouple probes.

(P) Changes in $T_{core}$ (black line) and $T_{BAT}$ (green line) following CNO administration with or without ice-cold water spray-induced $T_{core}$ at 33°C.

(Q) Experimental design for chemogenetic viral targeting of DMH Calb1$^-$/Vglut2$^+$ neurons in *Calb1$^{Cre}$* mice, with concurrent $T_{core}$ and EMG recordings in freely moving subjects.

(R) Representative synchronized cervical muscle EMG and $T_{core}$ traces corresponding to Figure 4B. Data were analyzed across the indicated dashed-line time intervals.

Data are presented as mean ± SD, n = 5 or 6 mice. Paired t-tests were used for statistical comparisons in panels B, C, F, J, K, and N.

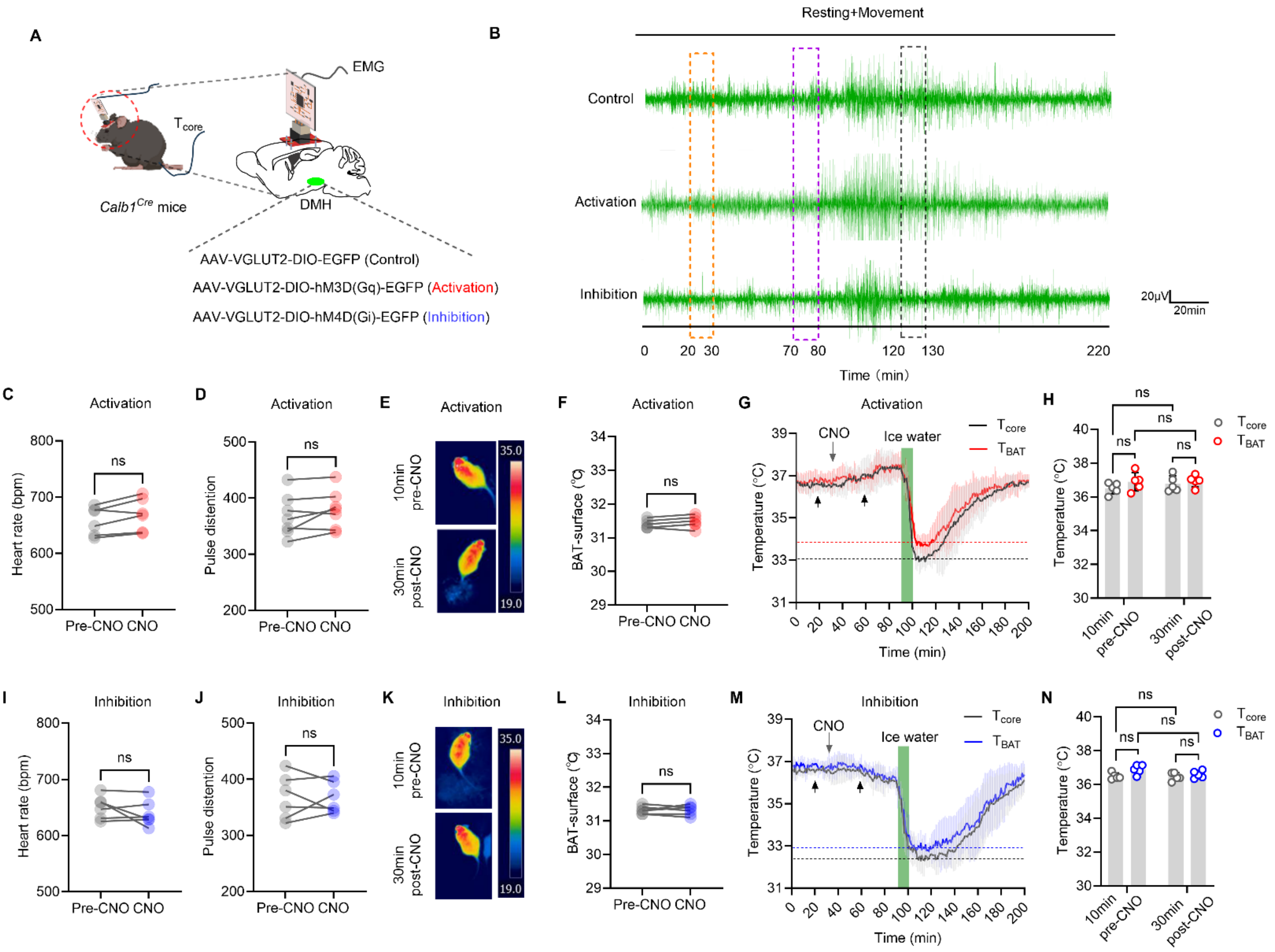


**Extended Data Fig. 5: Chemogenetic manipulation of DMH$^{Calb1/Vglut2}$ neurons selectively regulates hypothermia-induced shivering, with no effect on NST.**

(A) Schematic of chemogenetic viral injection into the DMH of *Calb1$^{Cre}$* mice; $T_{core}$ and EMG were recorded in freely behaving mice.

(B) Representative concurrent cervical muscle EMG and $T_{core}$ traces (aligned to Fig. 3B), sampled within dashed-line time windows.

(C, D, I, J) Quantified heart rate (C, I) and pulse distension (D, J) 10 min before and 30 min post-CNO injection, following chemogenetic activation (C, D) or inhibition (I, J) of DMH$^{Calb1/Vglut2}$ neurons in freely behaving mice.

(E, F, K, L) Representative infrared interscapular brown adipose tissue (iBAT) temperature ($T_{BAT}$) images (E, K) and corresponding quantification (F, L) pre-CNO and post-CNO for neuronal activation/inhibition groups.

(G, M) Simultaneous recordings of $T_{core}$ and $T_{BAT}$ of the iBAT using thermocouples in free-moving mice. Changes in $T_{core}$ (black lines) and $T_{BAT}$ (red or blue lines) after chemogenetic activation

(G) or inhibition (M) of $DMH^{Calb1/Vglut2}$ neurons with or without ice-cold water spray to reduce $T_{core}$ to 33°C.

(H, N) Quantitative analysis of $T_{core}$ and $T_{BAT}$ at 10 min prior to and 30 min following CNO administration in panels G and M (black arrows).

Data were presented as means ± SD, with n = 6 mice. Two-way ANOVA with Tukey's *post hoc* test was used for panels H, N, and Paired t test was used for panels C, D, F, I, J, L.

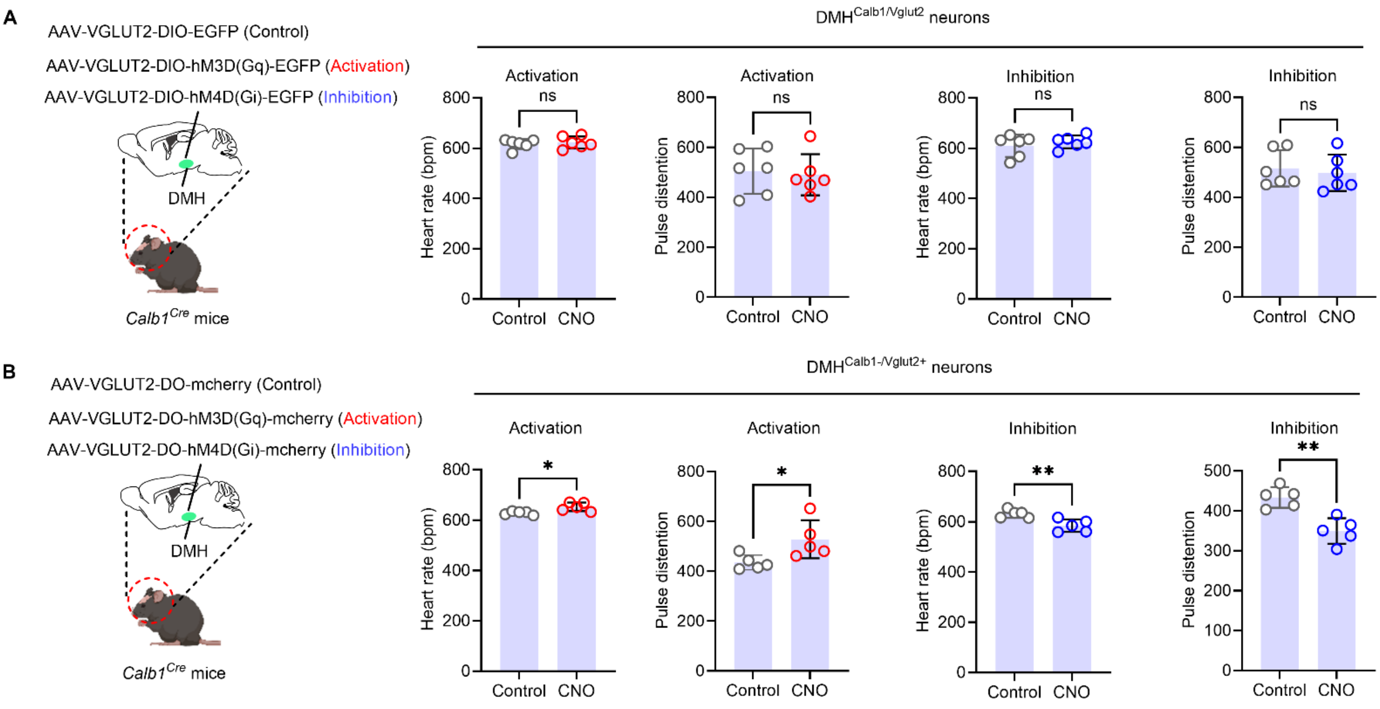


**Extended Data Fig. 6: Chemogenetic modulation of Calb1+/Vglut2+ and Calb1-/Vglut2+ neurons during hypothermia at $T_{core}$ 33°C.**

(A) Cre-on chemogenetic strategy targeting Calb1+/Vglut2+ neurons in the DMH. Left: schematic of viral delivery into the DMH of *Calb1$^{Cre}$* mice. Following cooling to 33 °C, chemogenetic activation or inhibition of DMH$^{Calb1/Vglut2}$ neurons, as well as measurements of heart rate and pulse distention, were conducted in freely moving mice.

(B) Cre-off chemogenetic strategy targeting Calb1–/Vglut2+ neurons in the DMH. Left: viral injection schematic in *Calb1$^{Cre}$* mice. Following cooling to 33 °C, chemogenetic activation or inhibition of DMH$^{Calb1\text{-}/Vglut2+}$ neurons, and heart rate and pulse distention, were measured and quantified in freely moving mice.

Data were presented as means ± SD, with n = 5 or 6 mice as indicated in the graphs. An unpaired t-test was used for statistical analysis. *$p < 0.05$, **$p < 0.01$.

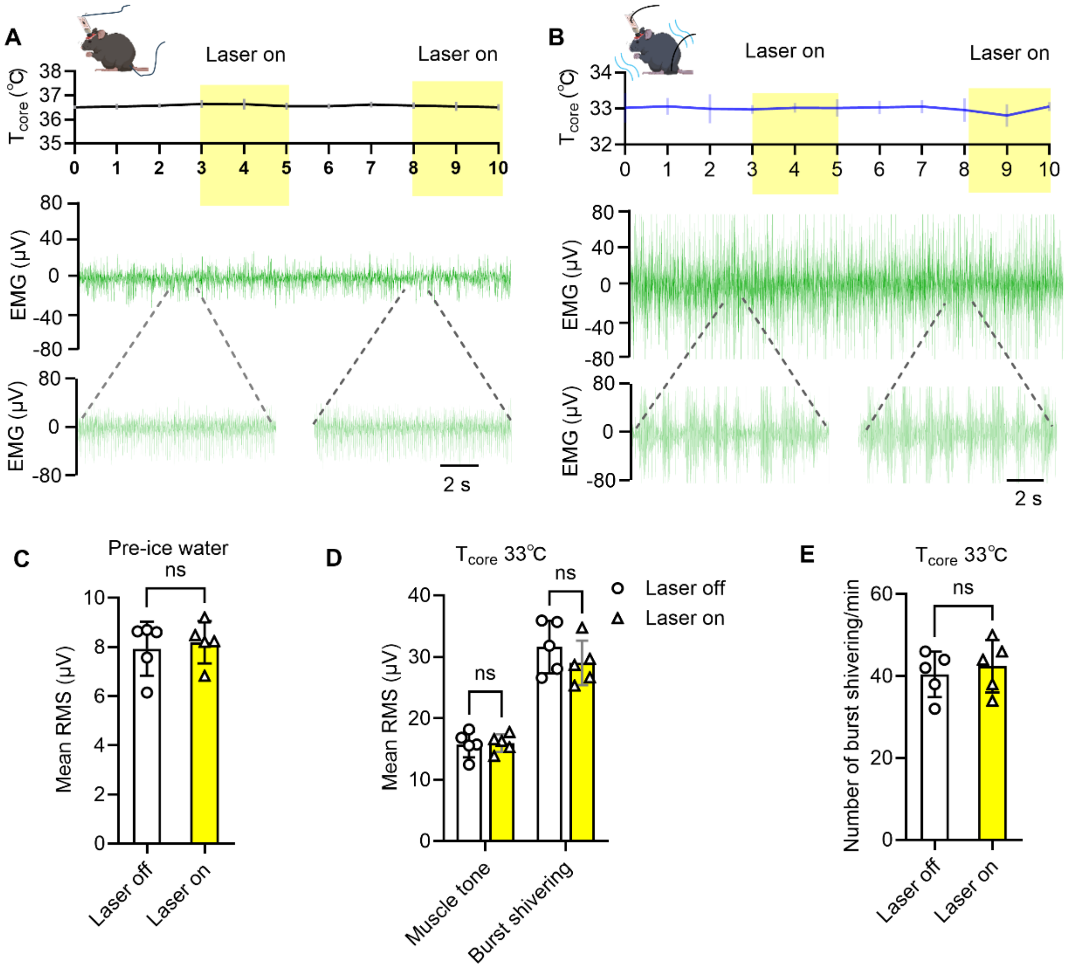


**Extended Data Fig. 7: Stimulation of the DMH with optogenetic control AAVs fails to alter cervical muscle EMG at normothermia and hypothermia.**

(A) Following injection of control virus (AAV2/9-VGLUT2-DIO-EGFP-WPREs) into the DMH of *Calb1*$^{Cre}$ mice, concurrent $T_{core}$ and cervical EMG were recorded during DMH opto-stimulation (laser-on: yellow shaded periods; laser-off: white periods) at normothermia ($T_{core}$ = 36.5 °C).

(B) Following delivery of control AAV2/9-VGLUT2-DIO-EGFP-WPREs to the DMH of *Calb1*$^{Cre}$ mice, $T_{core}$ and cervical EMG were recorded concurrently during DMH opto-stimulation (laser-on: yellow; laser-off: white) at hypothermic 33 °C.

(C) Quantification of resting EMG power comparing laser-on vs. laser-off states from panel A.

(D, E) Quantification of resting EMG power and shivering burst amplitude (D), plus shivering burst count (E) across laser states at 33 °C (panel B).

Data were presented as means ± SD, with n = 5 mice. A paired t-test was used for panels C and E. One-way ANOVA with Tukey's *post hoc* test was used for panel D.

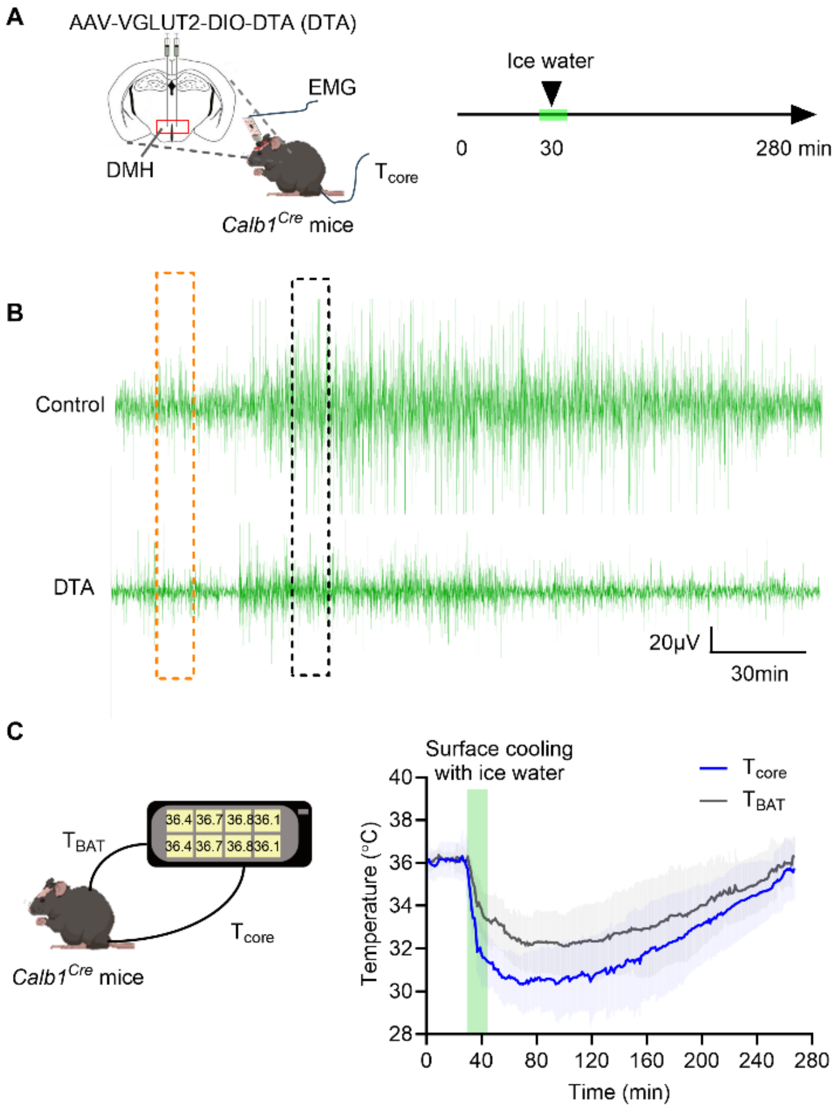


**Extended Data Fig. 8: Ablation of DMH$^{Calb1/Vglut2}$ neurons suppresses hypothermia-induced shivering without affecting hypothermia-induced $T_{BAT}$.**

(A) Schematic of DTA expression in the DMH of *Calb1$^{Cre}$* mice for targeted DMH$^{Calb1/Vglut2}$ neuronal ablation.

(B) Concurrent recordings of cervical EMG and $T_{core}$ in freely moving mice (corresponding to Fig. 5K). EMG power spectra within dashed-line time windows were analyzed quantitatively.

(C) Schematic of DTA viral injection into the DMH of *Calb1$^{Cre}$* mice to ablate DMH$^{Calb1/Vglut2}$ neurons. Simultaneous $T_{core}$ and $T_{BAT}$ recordings were performed via thermocouples in freely moving ablated mice during surface cooling.

Data were presented as means ± SD, with n = 6 mice.

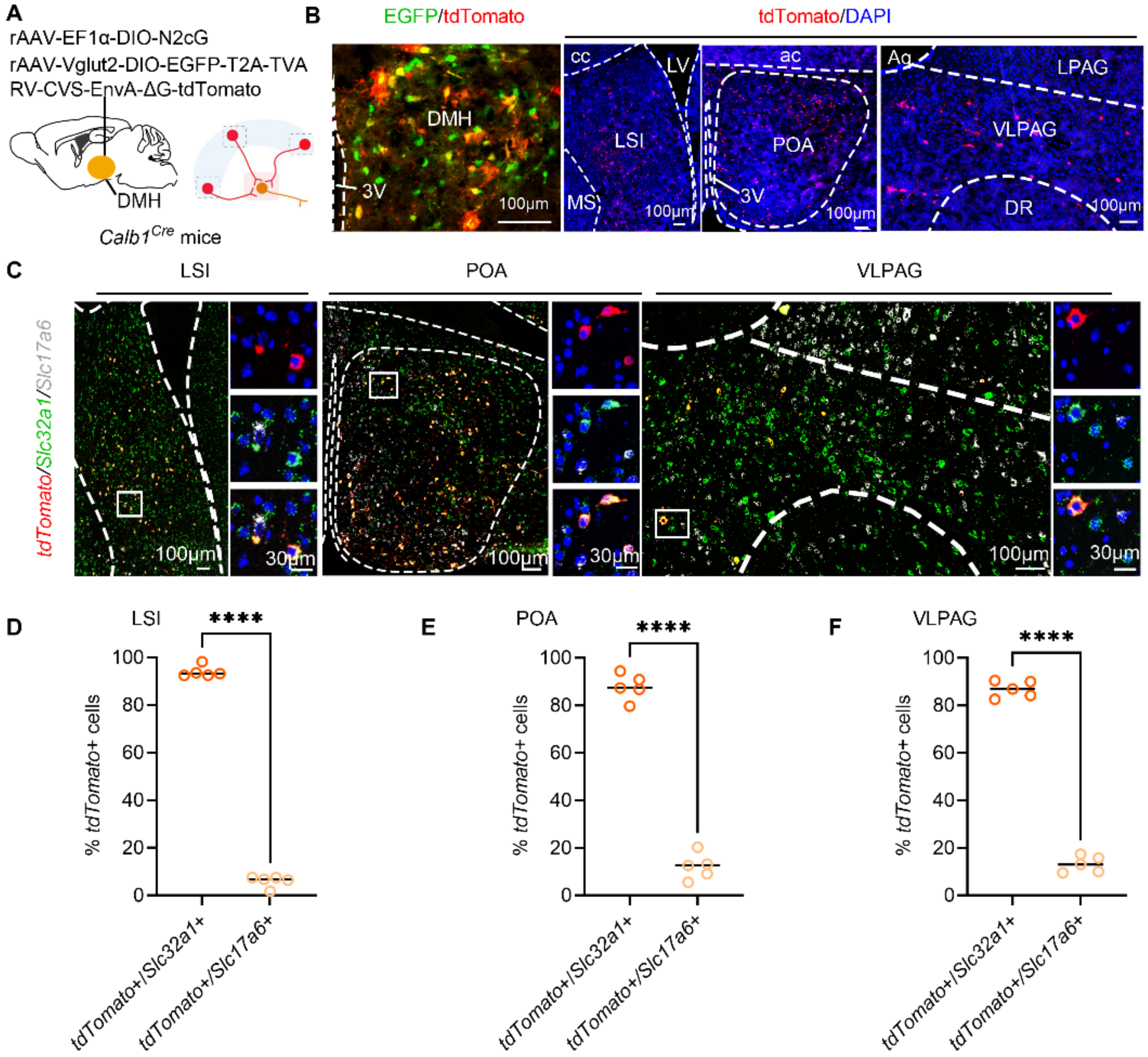


**Extended Data Fig. 9: DMH$^{Calb1/Vglut2}$ neurons receive predominantly inhibitory upstream innervation.**

(A) Schematic of retrograde viral tracing to map presynaptic inputs onto DMH$^{Calb1/Vglut2}$ neurons.

(B) Representative images showing robust tdTomato-labeled upstream neurons in the LSI, POA and VLPAG following monosynaptic retrograde tracing from EGFP/tdTomato double-positive DMH$^{Calb1/Vglut2}$ neurons.

(C) FISH staining for *Slc32a1* (VGAT, inhibitory) and *Slc17a6* (VGLUT2, excitatory) in *tdTomato*-positive input neurons; magnified view of the boxed region is shown; scale bars are as indicated on the micrographs.

(D–F) Quantification of the fractions of *tdTomato*+ input neurons expressing *Slc32a1* or *Slc17a6* from panel C.

Data were presented as means ± SD, with n = 5 mice. An unpaired t-test was used for statistical analysis. ****$p < 0.0001$.

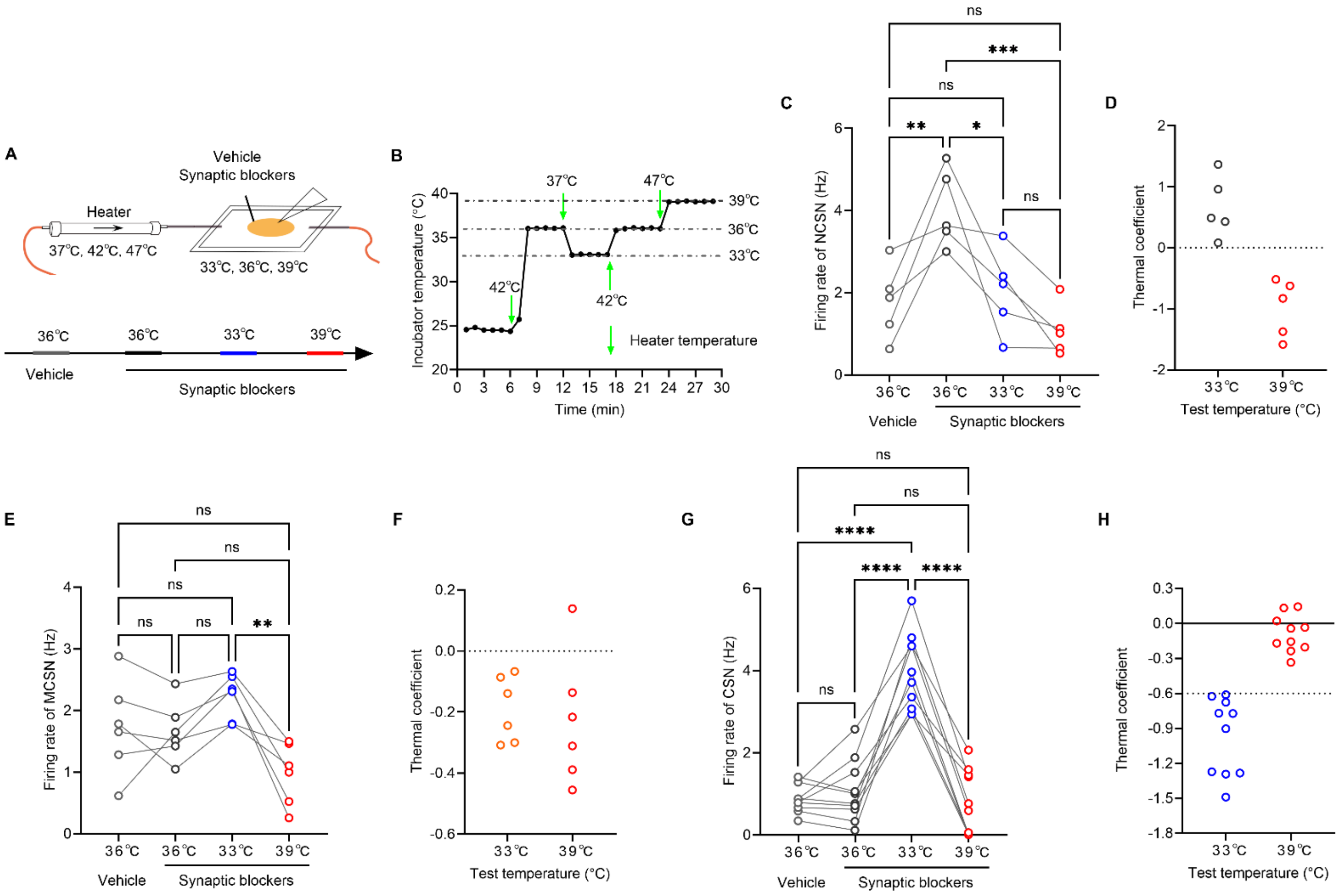


**Extended Data Fig. 10: DMH$^{Calb1/Vglut2}$ are not warm-sensitive neurons.**

(A) Schematic for whole-cell patch-clamp recordings of DMH$^{Calb1/Vglut2}$ neurons under temperature-controlled aCSF perfusion. Recordings were first obtained with a vehicle, then with a synaptic blocker cocktail (20 μM DNQX, 50 μM D-AP5, 20 μM gabazine) to isolate intrinsic action potential firing.

(B) Schematic of recording chamber temperature regulation at 33 °C, 36 °C and 39 °C. Inlet heater was adjusted to 37 °C, 42 °C and 47 °C, respectively, to stabilize bath temperature during recordings.

(C, E, G) Firing rate quantification for non-cold-sensitive (NCSN), moderately cold-sensitive (MCSN), and cold-sensitive neurons (CSN).

(D, F, H) Thermal coefficient (TC) analyses corresponding to panels C, E, G. Classification criteria (units: spikes·$s^{-1}$·°$C^{-1}$): warm-sensitive (TC > +0.8), NCSN (TC > 0), MCSN (−0.6 < TC < 0), CSN (TC ≤ −0.6).

Data were presented as means ± SEM, with n = 6 mice. One-way ANOVA with Tukey's *post hoc* test was used for statistical analysis. *p < 0.05, **p < 0.01, ***p < 0.001, ****p < 0.0001.

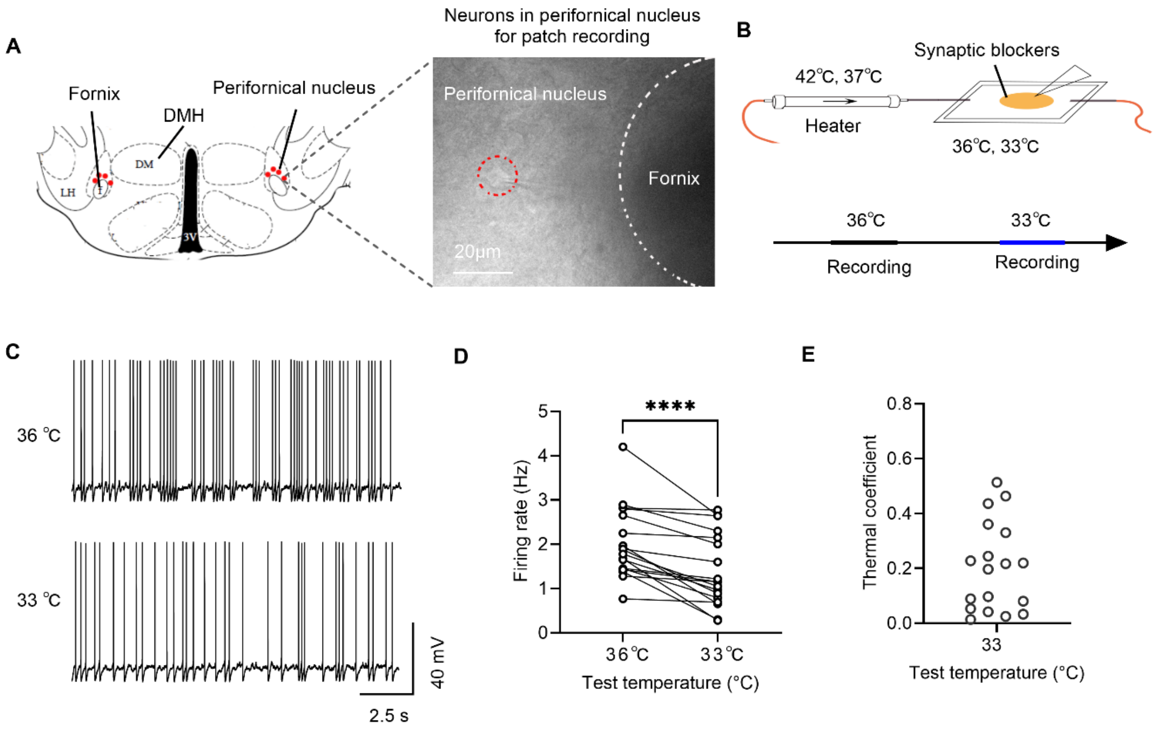


**Extended Data Fig. 11: Neurons in the perifornical nucleus adjacent to the DMH lack cold sensitivity.**

(A) Schematic and brain slice preparation showing patch-clamp-recorded neurons (red circles) in the DMH-adjacent perifornical nucleus.

(B) Schematic of temperature-controlled patch-clamp recording chamber with aCSF perfusion containing synaptic blockers (20 μM DNQX, 50 μM D-AP5, 20 μM gabazine).

(C) Representative current-clamp action potential traces of perifornical neurons recorded at 36 °C and 33 °C.

(D) Quantitative analysis of neuronal firing rates corresponding to panel C.

(E) Plot of perifornical neuron TC values which were all at dF/dT > 0 Hz/°C (n = 18 neurons), indicating failure to display a temperature-dependent rise in firing rate and did not satisfy the TC criterion characteristic of cold-sensitive neurons.

Data were presented as means, with n = 5 mice. A paired t-test was used for panel D. ****p < 0.0001.

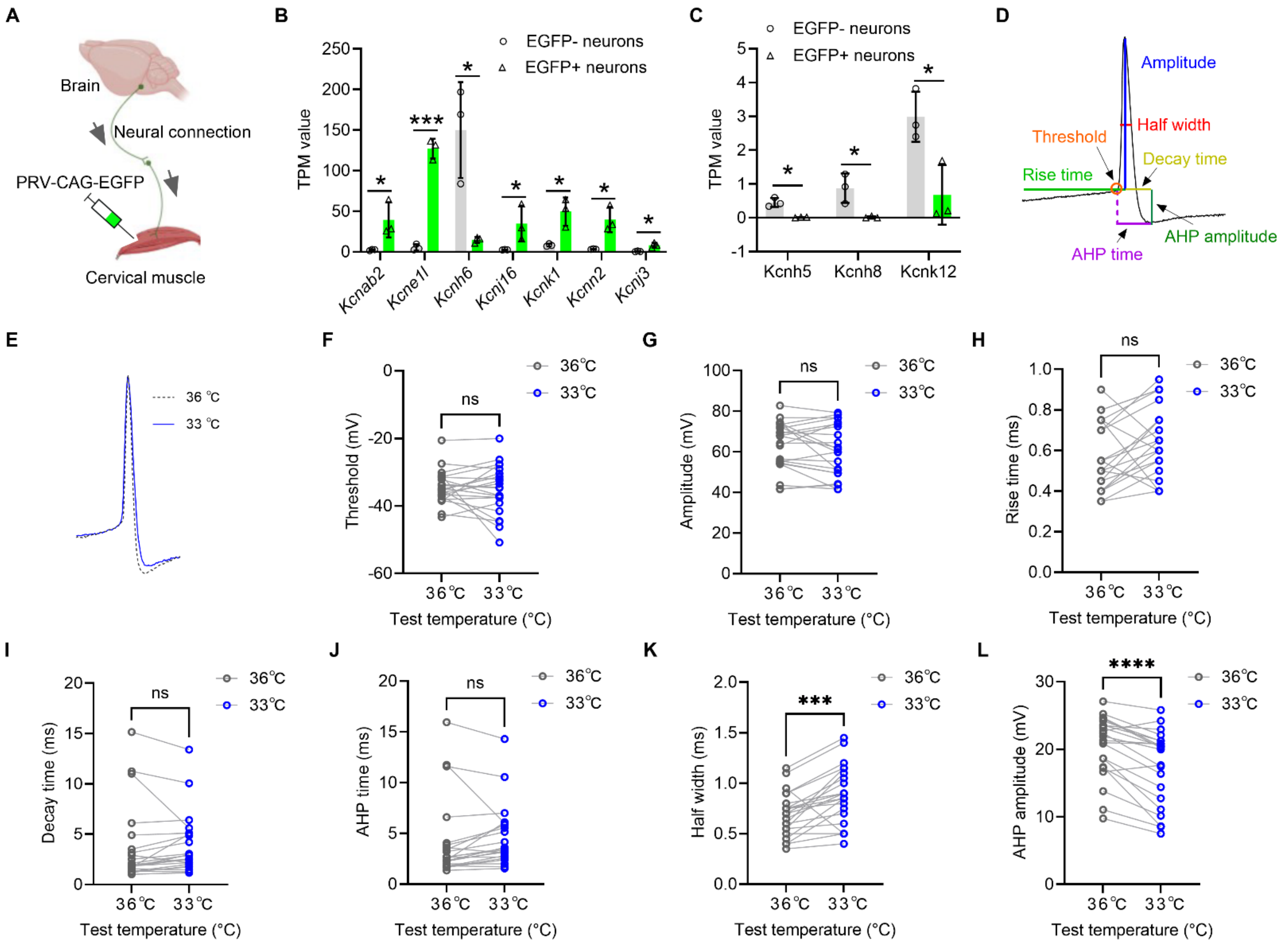


**Extended Data Fig. 12: Potassium channels potentially mediate hypothermia-induced activation of cold-sensitive DMH$^{Calb1/Vglut2}$ neurons.**

(A-C) Schematic of retrograde PRV-CAG-EGFP labeling of DMH neurons innervating cervical muscles (A). RNA-sequencing of EGFP+ and EGFP- neurons in DMH (see Supplemental Fig. 2A–D) revealed significant differences in potassium channel mRNA expression, quantified as transcripts per million (TPM) (B, C).

(D, E) Schematic action potential morphology (D) and representative recordings from CSN of DMH$^{Calb1/Vglut2}$ neurons (E).

(F-L) Quantitative analysis of key action potential properties: spike threshold (F), amplitude (G), rise time (H), decay time (I), afterhyperpolarization (AHP) duration (J), spike half-width (K), and AHP amplitude (L).

Data were presented as means, with n = 11 neurons in 5 mice. An unpaired t-test was used for panels B, C. A paired t-test was used for panels F-L. * $p < 0.05$, *** $p < 0.001$, **** $p < 0.0001$.
